# Human belief state-based exploration and exploitation in an information-selective symmetric reversal bandit task

**DOI:** 10.1101/2020.08.31.276139

**Authors:** Lilla Horvath, Stanley Colcombe, Michael Milham, Shruti Ray, Philipp Schwartenbeck, Dirk Ostwald

## Abstract

Humans often face sequential decision-making problems, in which information about the environmental reward structure is detached from rewards for a subset of actions. In the current exploratory study, we introduce an information-selective symmetric reversal bandit task to model such situations and obtained choice data on this task from 24 participants. To arbitrate between different decision-making strategies that participants may use on this task, we developed a set of probabilistic agent-based behavioral models, including exploitative and explorative Bayesian agents, as well as heuristic control agents. Upon validating the model and parameter recovery properties of our model set and summarizing the participants’ choice data in a descriptive way, we used a maximum likelihood approach to evaluate the participants’ choice data from the perspective of our model set. In brief, we provide quantitative evidence that participants employ a belief state-based hybrid explorative-exploitative strategy on the information-selective symmetric reversal bandit task, lending further support to the finding that humans are guided by their subjective uncertainty when solving exploration-exploitation dilemmas.

## Introduction

Uncertainty is an inherent part of real-life sequential decision making. Humans often face new and changing situations without being able to directly observe the statistical regularities of the environmental reward structure. Consequently, in their quest to maximize their cumulative rewards, humans have to alternate between exploration and exploitation. Exploration refers to decisions that maximize information gain and thus reduce the uncertainty about the statistical regularities of the environment. Exploitation refers to decisions that maximize reward gain by harnessing the accumulated knowledge.

A standard behavioral test bed to study sequential decision making under uncertainty is the bandit paradigm (e.g. Robbins, 1952; Brand et al., 1956; Brand & Woods, 1957; Berry & Fristedt, 1985; Cohen et al., 2007; Even-Dar et al., 2006; Dayan & Daw, 2008; Bubeck et al., 2009; Gabillon et al., 2012). Two variants of the bandit paradigm have been widely adopted to model real-life sequential decision making under uncertainty. We here refer to these variants as the *classical bandit paradigm* (by some also referred to as partial-feedback paradigm (c.f. Hertwig, 2012; Wulff et al., 2018)) and the *pure exploration paradigm* (by some also referred to as sampling paradigm, ibid.). In both variants, on each trial the deciding agent has to choose among a finite set of actions with different expected reward values and subsequently observes a reward with probability specific to the chosen action. While the actions’ expected rewards are not directly observable, the agent can estimate them by integrating information over multiple reward observations. The difference between the classical bandit paradigm and the pure exploration paradigm stems from the respective challenges they pose for the deciding agent. In the classical bandit paradigm, the agent’s task is to maximize the cumulative reward across all trials, while the reward observation confers both information and reward on each trial. The classical bandit paradigm thus raises the problem of how to strike a balance between exploration and exploitation on each trial. In contrast, in the pure exploration paradigm, the aim is to maximize the reward obtained on a single final trial. Here, reward observations on preceding trials only confer information, but not reward. The number of trials preceding this final trial is self-determined by the agent. The pure exploration paradigm thus raises the problem of how to strike a balance between the exploration costs preceding the final trial and the potential final trial reward (Ostwald et al., 2015).

While a large variety of real-life decision-making problems can be modelled with the classical bandit and the pure exploration paradigms, neither variant is suited to model a class of decision-making problems, in which each available action yields positive or negative reward, while only some actions also yield information about the problem’s reward structure. Consider for example the situation of a patient who exhibits COVID-19 symptoms during the global pandemic. Assuming that reliable and medically administered COVID-19 tests are available, while self-administered COVID-19 tests are not (as has been the case in many countries in the first year of the pandemic), the patient faces the following decision dilemma: on the one hand, the patient may home-quarantine, thereby minimizing the risk for virus transmission, but at the price of not obtaining information about their own state of infection. Alternatively, the patient may use public transport to undergo a medically administered COVID-19 test, thereby incurring the risk of further transmitting the virus, but obtaining reliable information about their personal infectious state. Situations of this type are similar to the ones modelled with the classical bandit paradigm as each action has a positively or negatively rewarded consequence (which in the example is of societal nature). Importantly, however, in this situation reward-relevant information is detached from reward for one of the actions (home-quarantining), akin to the pure exploration paradigm. Consequently, such a decision situation poses a more pronounced exploration-exploitation dilemma then both classical bandit paradigms and the pure exploration paradigm, because the decision maker is forced to explicitly evaluate the benefit of information gain against the benefit of reward gain. The aim of the current study is to computationally characterize human sequential decision making in such problems.

To this end, we introduce an information-selective symmetric reversal bandit task, which shares key characteristics with the classical symmetric two-armed reversal bandit task (e.g. Bartolo & Averbeck, 2020; Costa et al., 2016; Gläscher et al., 2009; Hauser et al., 2014), but in which information is randomly withheld for either the action with the high or the low expected reward value. To arbitrate between different sequential decision-making strategies that humans may employ on this task, we formulate a set of agent-based behavioral models. We here follow up on recent results showing that one way humans balance between exploration and exploitation is to add an ‘information bonus’ to the value estimate of an action, which reflects the associated uncertainty (e.g. Gershman, 2018, 2019; Lee et al., 2011; Wilson et al., 2014; Wu et al., 2018). More specifically, we formulate Bayesian agents that represent subjective uncertainty about the structure of the environment in the form of a belief state. The Bayesian agents use the belief state to make either exploitative (i.e., value estimate maximizing actions), explorative (i.e., information bonus maximizing actions), or hybrid explorative-exploitative (i.e., combined value estimate and information bonus maximizing) actions. Notably, we adopt a Bayesian treatment of exploration and quantify the information bonus as the expected Bayesian surprise (Itti & Baldi, 2009; Sun et al., 2011; Ostwald et al., 2012). In addition to the Bayesian agents, we also formulate belief state-free agents that implement simple strategies, such as a cognitive null model and the win-stay-lose-switch heuristic (Robbins, 1952). Upon validating our modeling initiative, we provide evidence for a belief state-based hybrid explorative-exploitative strategy based on choice data from 24 participants. In summary, we show that in a scenario where every decision has an economic consequence, but only some decisions are informative about the statistical reward structure of the environment, humans are guided by their subjective uncertainty when resolving the exploration-exploitation dilemma.

## Experimental methods

### Participants

Young adults were recruited from the Nathan Kline Institute Rockland Sample (NKI-RS), a community-ascertained and comprehensively characterized participant sample of more than 1000 individuals between 6 and 85 years of age (Nooner et al., 2012). We initially intended to enroll individuals from the lower and upper ends of the attention deficit hyperactivity disorder (ADHD) spectrum because we were interested in the relationship between ADHD symptoms and behavioral strategies in our task. Yet, the final sample of 24 individuals (12 female, 23 right-handed, age range: 18-35 years, mean age: 24.5 years, standard deviation age: 5.5 years) represented the mid-range of the ADHD spectrum. Moreover, individuals were only invited if they had no lifetime history of severe neurological or psychiatric disorder. We therefore treated the group of participants as a healthy sample and did not conduct analyses to relate ADHD symptoms to task behavior. For additional details about the recruitment and sample characteristics, please refer to Section S.1: Sample characteristics.

### Procedure

The study consisted of a one-time visit of 3.5 hours to the Nathan Kline Institute for Psychiatric Research (Orangeburg, NY, US). After providing written informed consent, participants were first requested to fill out a series of questionnaires measuring symptoms of ADHD and other mental disorders. Next, participants received detailed written instructions about the information-selective symmetric reversal bandit task and were encouraged to ask any clarification questions. For the detailed instructions provided to the participants, please refer to Section S.2: Participant instructions. To familiarize participants with the task, they next completed a test run of the task on a desktop computer. Finally, participants completed two experimental task runs in a Magnetic Resonance Imaging (MRI) scanner, while behavioral, eye tracking, and functional MRI data were acquired. Note that in the current work, we only report results from the analysis of the behavioral data acquired during MR scanning. The visit ended with the participants receiving a reimbursement of $100 (see below for details).

### Experimental design

We developed a symmetric two-armed reversal bandit task, in which the available actions were not only associated with varying expected reward values but also with varying information gains (information-selective symmetric reversal bandit task, Figure 1a). More specifically, on each task trial participants could decide between the actions of choosing a square on the right of the computer screen vs. choosing a triangle on the left of the screen, or between the actions of choosing a square on the left vs. choosing a triangle on the right of the screen. Depending on the shape chosen, the action was either lucrative and returned a reward of +1 with a probability of 0.85 and a reward of −1 with a probability of 0.15, or detrimental and returned a reward of +1 with a probability of 0.15 and a reward of −1 with a probability of 0.85. Depending on the side of the shape chosen, the action was also either informative and the returned reward was revealed to the participant, or it was non-informative and the returned reward was not revealed to the participant. Specifically, following an informative action either an image of a moneybag was displayed to signal a positive reward of +1, or an image of a crossed-out moneybag was displayed to signal the negative reward −1. In contrast, following a non-informative action an image of a question mark moneybag was displayed for both the rewards of +1 and −1. Importantly, while the actions’ lucrativeness was not directly observable and could only be inferred from the revealed rewards, the actions’ informativeness was directly observable throughout the experiment. In particular, for half of the participants the right screen side was associated with the informative action and the left screen side was associated with the non-informative action. For the other half of the participants the coupling between screen side and action informativeness was reversed. As a visual reminder for the participants, the informative and non-informative screen sides were also indicated by black and grey backgrounds, respectively. Note that we use the terms informative side and non-informative side in accordance with the action definitions. Similarly, we will also use the terms ‘lucrative shape’ and ‘detrimental shape’ instead of ‘action of choosing the lucrative or detrimental shape’ for simplicity. Also note that throughout the depiction of the experimental design and results, we visualize lucrative and detrimental actions by yellow and blue colors, respectively, and informative and non-informative actions by black and grey colors, respectively.

**Figure 1.**
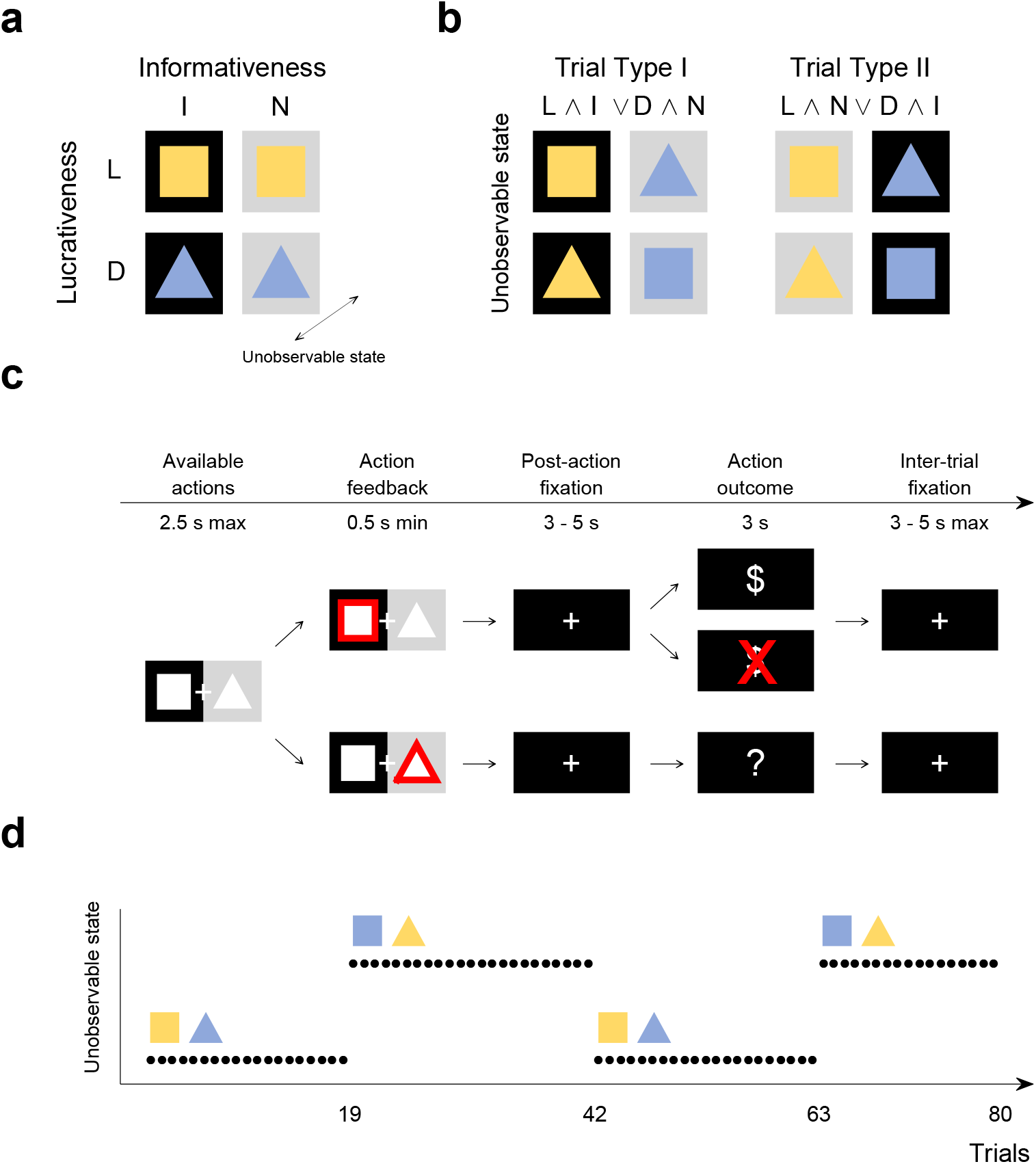
Information-selective symmetric reversal bandit task. **a** Experimental design. Possible actions differed in lucrativeness (lucrative (L) or detrimental (D)), as well as in informativeness (informative (I) or non-informative (N)). The former experimental factor was associated with selectable shapes (square and triangle), while the latter experimental factor was associated with black and grey screen sides. In addition, and unbeknownst to the participants, the lucrativeness of the shapes reversed at random trials throughout the experiment, corresponding to the unobservable task state: for some time, the square may represent the lucrative action, indicated here by its yellow color (and the triangle, accordingly, represent the detrimental action, indicated here by its blue color), but this could reverse. **b** On a given trial, participants faced a choice between either a lucrative and informative vs. a detrimental and non-informative action (Trial Type I; L ∧ I or D ∧ N) or a lucrative and non-informative vs. a detrimental and informative action (Trial Type II; L ∧ N or D ∧ I). **c** Trial design. Participants could indicate their choice within 2.5 seconds of the choice options onset. If they chose the shape on the black side, the returned reward was revealed (top). If they chose the shape on the grey side, the returned reward was not revealed (bottom). Note that the current lucrativeness of the shapes was not revealed to the participants, hence the white shape color. **d** Run design. Every 17 to 23 trials the reward probabilities associated with the shapes reversed. Here, the reversal times of the first run are shown. For trials 1 to 19 the square was lucrative, indicated by its yellow color, and the triangle was detrimental, indicated by its blue color. This reversed on trial 20 at which choosing the triangle became the lucrative action and choosing the square became the detrimental action. For the generation of this figure, please see *figure_1.m*.

The experiment consisted of two runs of 80 trials each. On half of the trials, choosing the square was lucrative and choosing the triangle was detrimental. On the other half of the trials, choosing the square was detrimental and choosing the triangle was lucrative. We pseudorandomized the sequence of lucrative shapes, such that choosing a certain shape was lucrative for 17-23 consecutive trials upon which the actions’ lucrativeness reversed. This yielded a total of three shape lucrativeness reversals (or equivalently, four blocks of trials without a reversal) per task run (Figure 1d). Furthermore, we also pseudo-randomized the trial-by-trial sequence of choice options (e.g., a choice between the square on the informative side or the triangle on the non-informative side) with two constraints. First, a certain choice option combination occurred for a maximum of five consecutive trials. Second, on 50% of the trials in which the square was lucrative, the square was presented on the informative side (and the triangle on the non-informative side), while on the other 50% of the trials, the square was presented on the non-informative side (and the triangle on the informative side). The same constraint applied to those trials on which the triangle was lucrative. This way we did not only counterbalance the shape-side combinations, but also ensured that participants faced a choice between a lucrative and informative action (L ∧ I) and a detrimental and non-informative (D ∧ N) action on half of the trials. We refer to the trials with action choices between L ∧ I and D ∧ N actions as Trial Type I (Figure 1b). Accordingly, on the other half of the trials, participants faced a choice between a lucrative and non-informative action (L ∧ N) and a detrimental and informative (D ∧ I) action. We refer to the trials with action choices between L ∧ N and D ∧ I actions as Trial Type II (Figure 1b). Importantly, for a consistent experiment history across participants, we generated the sequence of lucrative shapes and choice options prior to the study and used the identical trial sequence for all participants. The task was implemented as *irb_task.py* in Python 2.7 using PsychoPy V1.82.01 (Peirce, 2007).

Participants were encouraged to maximize the cumulative sum of returned rewards across all trials. As an incentive, participants were informed that in addition to a standard reimbursement of $70 for partaking in the study, they would receive a bonus up to $30 depending on their final balance at the end of the second run of the task. They were not further informed about the balance-bonus conversion rate. In effect, however, all participants were payed the full bonus of $30 as requested by the Institutional Review Board.

### Trial design

Each trial started with the presentation of the two available choice options and participants were given a maximum of 2.5 seconds to indicate their choice (Figure 1c). If participants responded within this time window, the border of the chosen shape turned white to signal the recording of their choice. The duration of this feedback signal depended on the response time, such that the choice options and feedback together were presented for 3 seconds in total. Then, a post-choice fixation cross was presented for 3-5 seconds. This fixation cross was followed by the image representing the choice outcome, i.e., a moneybag, a crossed-out moneybag, or a question mark moneybag image, which was presented for 3 seconds. Finally, before the start of the next trial, an inter-trial fixation cross was displayed for 3-5 seconds. If participants did not respond within the choice time window, the message ‘too slow’ appeared for 0.5 seconds followed by an inter-trial fixation cross, a reward of −1 was automatically registered to the participant’s account, and the next trial commenced. Notably, while the sequences of lucrative shapes and choice options were generated prior to the experiment, the fixation cross duration times and the returned rewards were sampled online as participants interacted with the task. Specifically, the fixation cross duration times were sampled uniformly from an interval of 3 to 5 seconds. The reward values +1 and −1 were sampled from discrete categorical distributions with probabilities 0.85 and 0.15 for the lucrative action and with probabilities 0.15 and 0.85 for the detrimental action, respectively.

## Descriptive analyses

To assess the behavioral data set on a descriptive level, we evaluated nine summary choice rates for every participant. In particular, we first evaluated overall and trial type-specific valid choice rates. These choice rates were defined as the number of valid action choices on all trials, on Type I trials, and on Type II trials divided by the number of all trials, of Type I trials, and of Type II trials, respectively. For example, by design there were 80 trials of Type I. If a participant failed to make a valid choice on one of these trials, the Trial Type I valid choice rate evaluated to 79/80. The participant-specific choice rates were then averaged across participants and the standard error of the mean (SEM) was evaluated. These analyses showed that participants completed the vast majority of trials and achieved an overall valid choice rate of 97.86% ± 0.62. There was virtually no difference in the number of valid choices between trial types: The valid choice rate on Trial Type I was 97.97% ± 0.62 and the valid choice rate on Trial Type II was 97.76% ± 0.68.

We then evaluated the choice rates for the lucrative and informative actions (L ∧ I), lucrative and non-informative actions (L ∧ N), detrimental and informative actions (D ∧ I), and detrimental and non-informative actions (D ∧ N). These choice rates were computed by dividing the number of respective actions by the number of valid choices of the corresponding trial type. Consequently, the choice rates of a given trial type are symmetrical, i.e., they sum up to 100%. For example, if a participant on Type I trials made 79 valid action choices of which 65 were L ∧ I actions and 14 were D ∧ N actions, then the L ∧ I choice rate was 65/79 and the D ∧ N choice rate was 14/79. In addition, we evaluated the choice rates of the lucrative actions and the informative actions. These were computed by dividing the sum of the number of L ∧ I and L ∧ N actions, as well as the sum of the number of L ∧ I and D ∧ I actions, by the number of valid choices on all trials. For example, if a participant made 159 valid choices in total, and of these 65 choices were L ∧ I actions, while 58 were L ∧ N actions, the lucrative action choice rate evaluated to 123/159. The participant-specific choice rates were then averaged across participants and SEM was evaluated. As shown in Figure 2a, on Trial Type I, the majority of action choices was lucrative and informative (L ∧ I, 87.45%± 1.53), while only a few action choices were detrimental and non-informative (D ∧ N, 12.55%±1.53). The difference between the choice rates on Trial Type II was less pronounced: as shown in Figure 2a, 66.01% ± 2.28 of the action choices on Trial Type II were lucrative and non-informative (L ∧ N), while 33.99% ± 2.28 were detrimental and informative (D ∧ I). Summed over informative and non-informative action choices, the lucrative action choice rate was 76.74% ± 1.7, whereas summed over lucrative and detrimental action choices, the informative action choice rate was 60.74% ± 0.92. Notably, participants made more lucrative choices if the lucrative action was also informative (L ∧ I, 87.45% ± 1.53) compared to lucrative choices if the lucrative action was non-informative (L ∧ N, 66.01%± 2.28). To statistically corroborate this finding, we conducted a two-sided paired sample t-test across participants. This yielded a test statistic of *t*(23) = 11.55 with an associated p-value smaller than 0.001. Taken together, these summary statistics suggest that while participants’ choices were primarily guided by action lucrativeness, participants also took the action’s informativeness into account when deliberating which action to choose.

**Figure 2.**
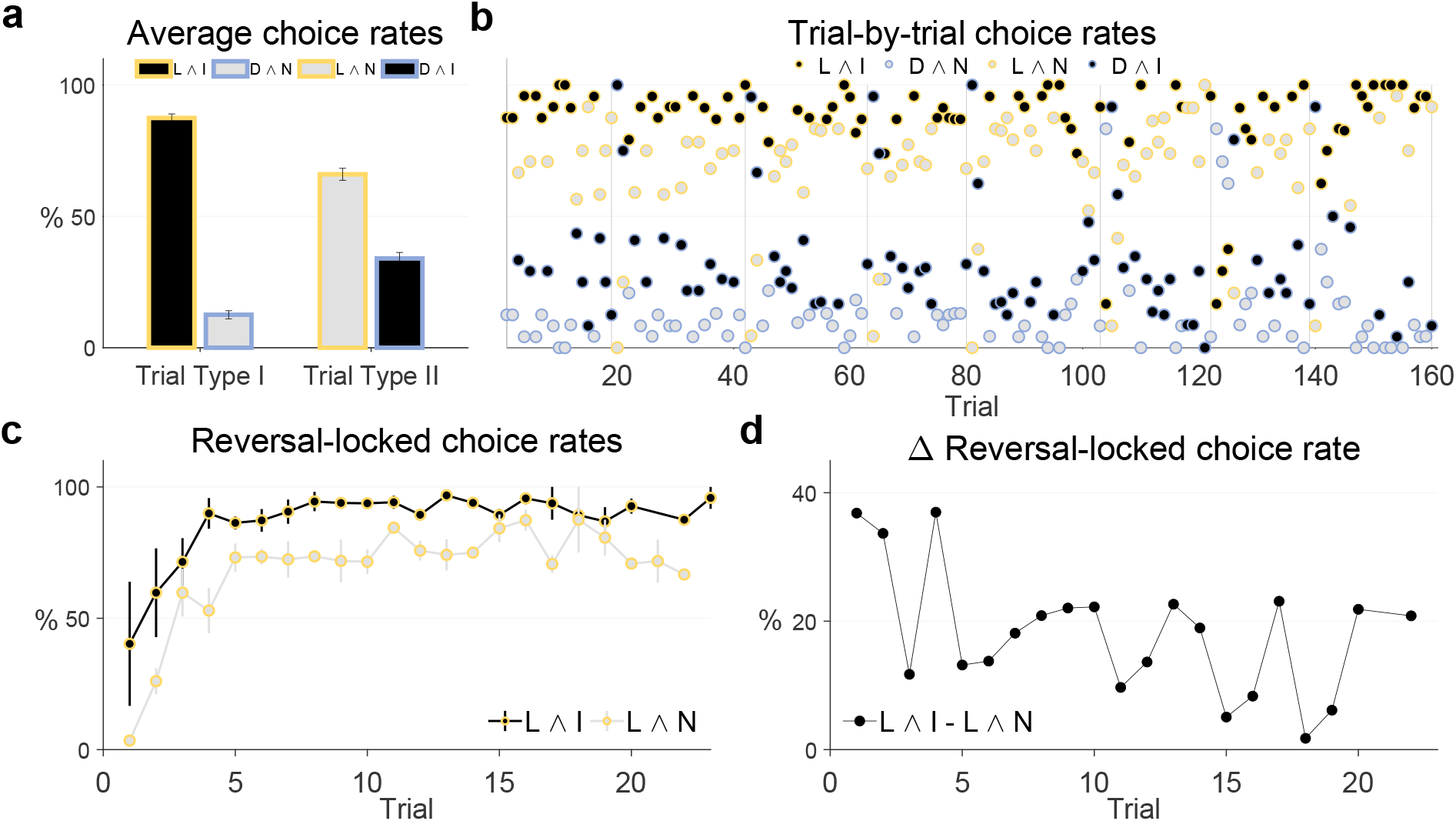
Participant choice rates. **a** Average choice rates across participants and task dynamics for Trial Type I and Trial Type II. On Trial Type I, participants faced a choice between a lucrative and informative (L ∧ I) and a detrimental and non-informative (D ∧ N) action. On average, participants preferred the L ∧ I action. On Trial Type II, participants faced a choice between a lucrative and non-informative (L ∧ N) and a detrimental and informative (D ∧ I) action. On average, participants preferred the L ∧ N action, but to a lesser degree than the L ∧ I action on Trial Type I. **b** Group trial-by-trial L ∧ I, D ∧ N, L ∧ N, and D ∧ I action choice rates. The vertical lines represent the last trial before a reward rate reversal. The vertical lines at *t* = 80 and *t* = 160 mark the end of the first and second run, respectively. Note that given the experimental design, the action choice rates on Trial Type I (L ∧ I vs. D ∧ N) and on Trial Type II (L ∧ N vs. D ∧ I) are complementary and add up to 100%. On the majority of trials the lucrative action choice rates L ∧ I and L ∧ N prevail over the detrimental action choice rates D ∧ I and D ∧ N. This effect is more pronounced for the L ∧ I choice rate than for the L ∧ N choice rate. **c** Average reversal-locked group trial-by-trial action choice rates for L ∧ I and L ∧ N actions. Both action choice rates increase over post-reversal trials. The error bars depict the SEM over reversal blocks. **d** Average group reversal-locked L ∧ I and L ∧ N choice rate difference. The difference decreased between the first trials after and the last trials before a reversal. For implementational details, please see *figure_2.m*.

In addition to the summary choice rates, we also evaluated trial-by-trial choice rates. Specifically, we computed group trial-by-trial L ∧ I, L ∧ N, D ∧ I, and D ∧ N action choice rates. To this end, for every trial we divided the number of respective actions by the number of valid choices on the trial over participants. As a given trial belonged to one of the two trial types, it either had associated L ∧ I and D ∧ N action choice rates, or associated L ∧ N and D ∧ I action choice rates. Consequently, in accordance with the summary action choice rates, the choice rates of each trial were symmetrical. For example, by design the first trial of the first run was of Type I for every participant. If on this trial 18 participants chose the L ∧ I action, 5 chose the D ∧ N action, and 1 participant missed to make a valid choice, the L ∧ I action choice rate for this trial evaluated to 18/23 and the D ∧ N action choice rate evaluated to 5/23. Finally, for each trial between two reversals, we computed the average reversal-locked group trial-by-trial L ∧ I action and L ∧ N action choice rates. Note that because the trial sequence was pseudo-randomized, the average reversal-locked group choice rate of a particular trial was computed based on different number of data points. For example, of the eight first trials, three were of Type I and had an associated group trial-by-trial L ∧ I action choice rate, while five were of Type II and had an associated group trial-by-trial L ∧ N action choice rate. Also note that as the number of trials between two reversals varied, there were fewer than eight 18th to 23rd trials. As show in Figure 2b, on the majority of trials, the two lucrative trial-by-trial action choice rates L ∧ I and L ∧ N prevailed over the two detrimental trial-by-trial action choice rates D ∧ I and D ∧ N. This effect was more pronounced for the trial-by-trial L ∧ I action choice rate. Notably, as shown in Figure 2c, both the average L ∧ I action choice rate as well as the average L ∧ N action choice rate exhibit an overall increase between two reward rate reversals, indicating that participants were able to gradually resolve their uncertainty about the currently lucrative shape. Moreover, as shown in Figure 2d, although the average L ∧ I action choice rate was larger than the average L ∧ N action choice rate on virtually all trials between two reversals, their difference decreased slightly between the first trial after and the last trial before a reward rate reversal. This suggests that with decreasing uncertainty about the currently lucrative shape participants placed less valence on the actions’ informativeness.

## Agent-based behavioral modeling

To arbitrate between different trial-by-trial decision-making strategies that participants may have used on the experimental task and that gave rise to the descriptive results documented above, we used an agent-based behavioral modeling approach. In our documentation of this approach, we proceed as follows. In the *Model formulation* section, we first formulate the relevant model components, comprising a *task model*, a set of *agent models*, and a set of *data analysis models*. Here, the *task model* corresponds to a probabilistic model that captures key aspects of the experiment and serves to explicate the agents’ knowledge about their choice environment. The *agent models* specify the dynamic subjective representation of the task (e.g., trial-by-trial belief state updates for some agent models) and several decision-making processes based on these representations. Finally, the *data analysis models* specify the embedding of the agent models in a statistical observation framework, allowing for the quantification of decision noise and the estimation of the models’ parameters and evidence. Having formulated our modeling approach, we then document the computational methods for model parameter estimation and model evidence evaluation in the *Model estimation and comparison* section. Finally, we report the results of a number of model validation analyses (*Model and parameter recovery analyses*) and conclude with the evaluation of the agent-based behavioral models in the light of the experimental data (*Model comparison results*).

### Model formulation

#### Task model

To render the task amenable to agent-based behavioral modeling, we first formulated a model of the task using concepts from the theory of partially observable Markov decision processes (Bertsekas, 2000). Specifically, we represent an experimental run by the tuple

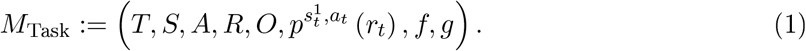

Here,

- *T* denotes the number of trials, indexed by *t* = 1, …, *T*.
- *S* ≔ ℕ_2_ × ℕ_2_ denotes the set of states *s* ≔ *s*^1^, *s*^2^. The first state component *s*^1^ encodes the lucrative shape. Specifically, on trial *t*, 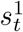 takes on the value 1 if the square is lucrative and takes on the value 2 if the triangle is lucrative. From the perspective of the agent, *s*^1^ is not directly observable. The second state component *s*^2^ encodes the available actions. Specifically, on trial *t*, 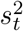 takes on the value 1, if the agent can choose between the square on the informative side or the triangle on the non-informative side. If on trial *t* the agent can choose between the square on the non-informative side or the triangle on the informative side, 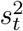 takes on the value 2. From the perspective of the agent, *s*^2^ is directly observable.
- *A* ≔ {*A*_1_, *A*_2_} denotes the set of state-dependent action sets. Specifically, depending on the observable state component 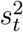 on a given trial *t* the available actions are either *A*_1_ ≔ {1, 4} or *A*_2_ ≔ {2, 3} for 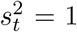 or 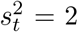, respectively. If the available action set is *A*_1_, then the agent can choose between *a* = 1, which corresponds to choosing the square on the informative side vs. *a* = 4, which corresponds to choosing the triangle on the non-informative side. If the available action set is *A*_2_, then the agent can choose between *a* = 2, which corresponds to choosing the square on the non-informative side vs. *a* = 3, which corresponds to choosing the triangle on the informative side.
- *R* ≔ {−1, +1} denotes the set of rewards *r*.
- *O* ≔ ℕ_3_ denotes the set of observations *o*. *o* = 1 encodes the image of the crossed-out moneybag, *o* = 2 encodes the image of the moneybag, and *o* = 3 encodes the image of the question mark moneybag.
- 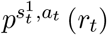 is the state- and action-dependent reward distribution. For each combination of *s*^1^ ∈ *S*^1^ and 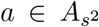, the state- and action-dependent reward distribution conforms to a discrete categorical distribution over *r_t_* with probability parameters listed in the first panel of Table 1. As an example, consider *s*^1^ = 1 (square is lucrative) and *a* = 1 (square on the informative side chosen). In this case, a reward of −1 is returned with a probability of 0.15 and a reward of +1 is returned with a probability of 0.85. On the other hand, if *s*^1^ = 2 (triangle is lucrative) and *a* = 1 (square on the informative side chosen), the reward probabilities are reversed.
- *f* is the state evolution function, which specifies the value the state *s_t_* takes on at trial *t*,

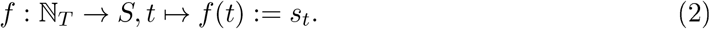

*f* is defined in a tabular form and corresponds to the sequence of lucrative shapes and choice options presented to all participants (cf. Section S.3: Experimental state sequence).
- *g* is the observation function

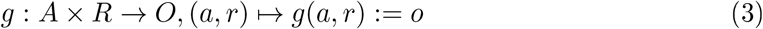

as defined in the second panel of Table 1. For the informative actions *a* = 1 and *a* = 3, *g* is injective: The reward *r* = −1 is mapped onto the observation *o* = 1, corresponding to the image of the crossed-out moneybag, while the reward *r* = +1 is mapped onto the observation *o* = 2, corresponding to the image of the moneybag. For the non-informative actions *a* = 2 and *a* = 4, *g* is not injective: both rewards *r* = −1 and *r* = +1 are mapped onto the observation *o* = 3, corresponding to the image of the question mark moneybag.

**Table 1.**
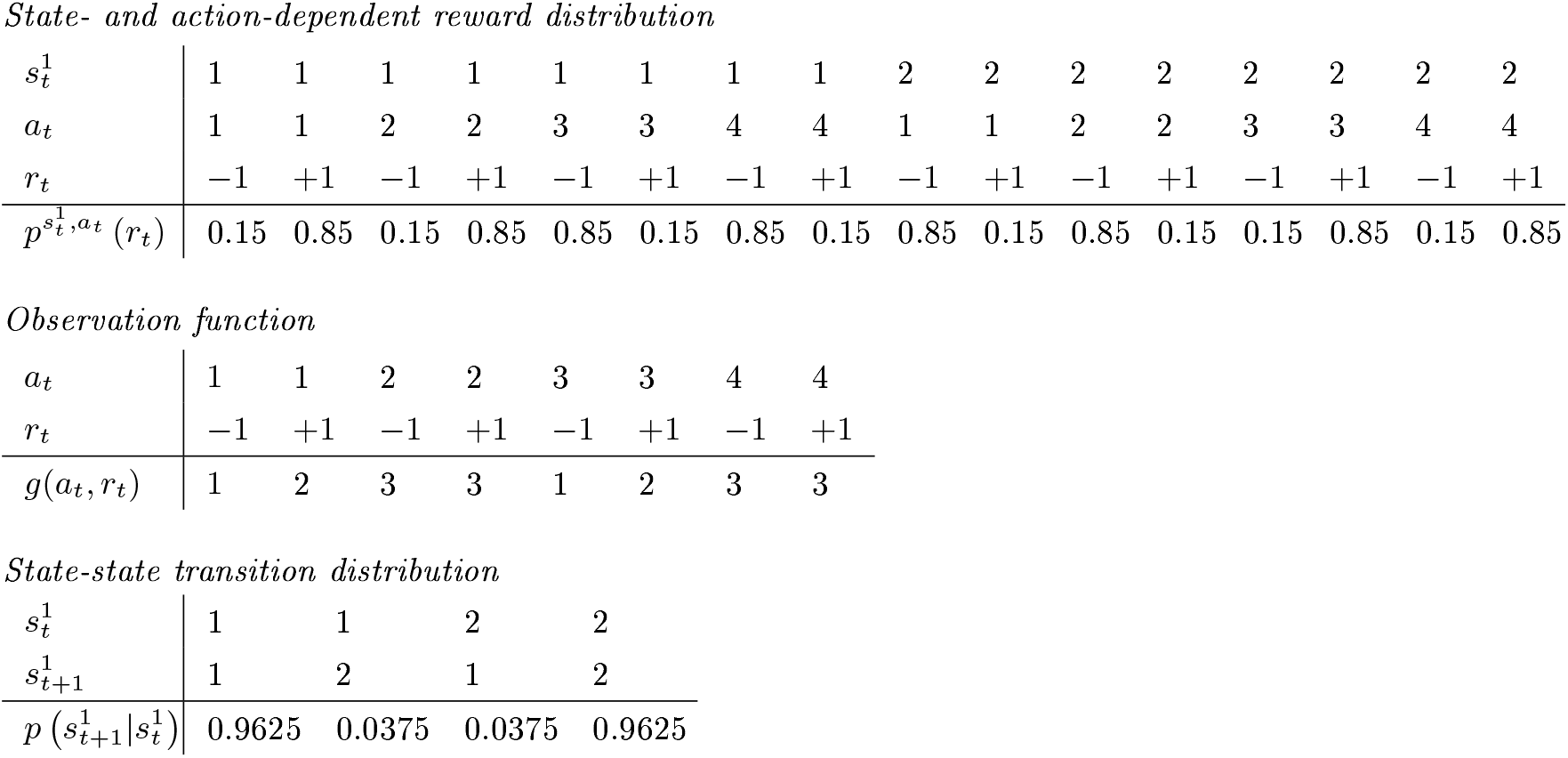
Model components. The upper table shows the state- and action-dependent reward distribution 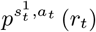, the middle table shows the observation function *g* and the lower table shows the action-independent state-state transition distribution 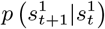.

|  |  |  |  |  |  |  |  |  |  |  |  |  |  |  |  |  |
| --- | --- | --- | --- | --- | --- | --- | --- | --- | --- | --- | --- | --- | --- | --- | --- | --- |
| $s_t^1$ | 1 | 1 | 1 | 1 | 1 | 1 | 1 | 1 | 2 | 2 | 2 | 2 | 2 | 2 | 2 | 2 |
| $a_t$ | 1 | 1 | 2 | 2 | 3 | 3 | 4 | 4 | 1 | 1 | 2 | 2 | 3 | 3 | 4 | 4 |
| $r_t$ | -1 | +1 | -1 | +1 | -1 | +1 | -1 | +1 | -1 | +1 | -1 | +1 | -1 | +1 | -1 | +1 |
| $p^{s_t^1, a_t}(r_t)$ | 0.15 | 0.85 | 0.15 | 0.85 | 0.85 | 0.15 | 0.85 | 0.15 | 0.85 | 0.15 | 0.85 | 0.15 | 0.15 | 0.85 | 0.15 | 0.85 |

Observation function
|  |  |  |  |  |  |  |  |  |
| --- | --- | --- | --- | --- | --- | --- | --- | --- |
| $a_t$ | 1 | 1 | 2 | 2 | 3 | 3 | 4 | 4 |
| $r_t$ | -1 | +1 | -1 | +1 | -1 | +1 | -1 | +1 |
| $g(a_t, r_t)$ | 1 | 2 | 3 | 3 | 1 | 2 | 3 | 3 |

**Table 1. Model components.** The upper table shows the state- and action-dependent reward distribution $p^{s_t^1, a_t}(r_t)$ , the middle table shows the observation function $g$ and the lower table shows the action-independent state-state transition distribution $p(s_{t+1}^1 | s_t^1)$ .
|  |  |  |  |  |
| --- | --- | --- | --- | --- |
| $s_t^1$ | 1 | 1 | 2 | 2 |
| $s_{t+1}^1$ | 1 | 2 | 1 | 2 |
| $p(s_{t+1}^1 \vert s_t^1)$ | 0.9625 | 0.0375 | 0.0375 | 0.9625 |

#### Agent models

We designed five agent models, denoted by C1, C2, A1, A2, and A3, to account for the putative cognitive processes underlying participants’ choices (cf. Table 2). Before we introduce the individual characteristics of these agents, we first represent the general structure of an agent interacting with an experimental run. This general agent structure corresponds to the tuple

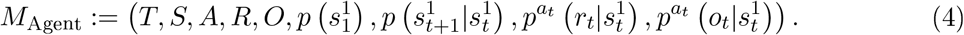

**Table 2.**
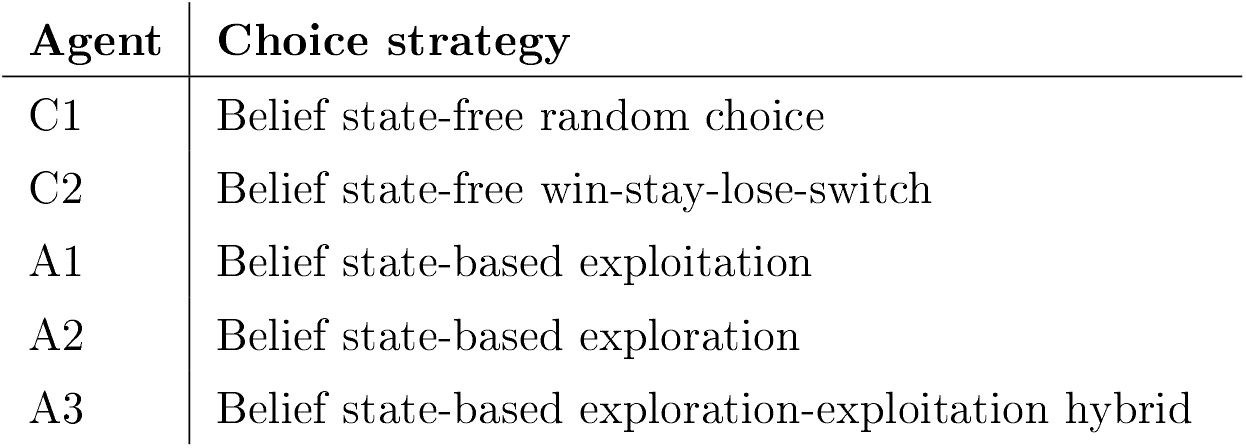
Agent model space. Agent model denominations (left column) and keywords highlighting central aspects of the respective agent’s choice strategy (right column).

Here,

- *T*, *S*, *A*, *R* and *O* are defined as the corresponding sets of the task model *M*_Task_.
- 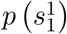 denotes the initial agent belief state, which specifies the agent’s subjective uncertainty over the non-observable state component 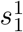 at trial *t* = 1. 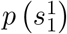 is defined in terms of the discrete categorical distribution

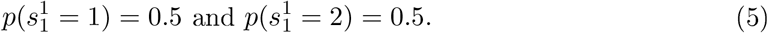

Because 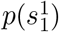 is fully parameterized by specifying 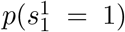, we hereinafter occasionally represent the initial belief state by the scalar 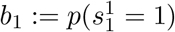.
- 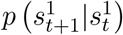 is the state-state transition distribution, which specifies the agent’s subjective uncertainty over the non-observable state component 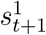 at trial *t* + 1 given the non-observable state component *s*^1^ at trial *t*. More specifically, for each *s*^1^ ∈ *S*^1^, the state-state transition distribution corresponds to a discrete categorical distribution over 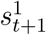 with probability parameters listed in the third panel of Table 1. Note that the trial-by-trial state transitions are probabilistic, because from the agent’s perspective a reversal in the shapes’ lucrativeness could happen between any two trials. This is in contrast with the state evolution from the task perspective, which - given the apriori defined sequence of lucrative shapes - is deterministic (cf. eq. (2)). Crucially, participants were informed that a reversal would happen 1-4 times in a run, but were not informed about the approximate number of trials without a reversal. Therefore, we equipped the agent with a constant reversal probability of 0.0375, which reflects the true reversal frequency in a run (there were 3 reversals across the 80 trials). For example, if 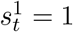 (square is lucrative), the agent allocates a probability of 0.9625 to the event that on the next trial 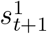 takes on the value 1 (square is lucrative), while it allocates a probability of 0.0375 to the event that 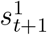 takes on the value 2 (triangle is lucrative).
- 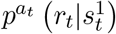 is the action-dependent state-conditional reward distribution, which specifies the agent’s subjective uncertainty over the reward *r_t_* given the non-observable state component *s*^1^ and action *a* at trial *t*. More specifically, for each combination of *s*^1^ ∈ *S*^1^ and 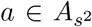, the action-dependent state-conditional reward distribution defines a discrete categorical distribution over *r_t_* with probability parameters corresponding to

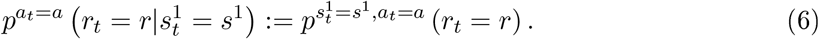

Notice that the only difference between the agent’s action-dependent state-conditional reward distribution and the task’s state- and action-dependent reward distribution is that for the former, the state is conceived as a random variable, while for the latter the state is conceived as a parameter. We equipped the agent with the true reward emission probabilities to reflect the task instructions. In particular, participants were truthfully informed that choosing the lucrative shape would return a reward of +1 with a high probability and a reward of −1 with a low probability, as well as that choosing the detrimental shape would return a reward of +1 with a low probability and a reward of −1 with a high probability.
- 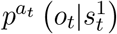 is the action-dependent state-conditional observation distribution, which specifies the agent’s subjective uncertainty over the observation *o_t_* given the non-observable state component *s*^1^ and action *a* at trial *t*. In detail, for each combination of *s*^1^ ∈ *S*^1^ and 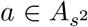, the action-dependent state-conditional observation distribution corresponds to a discrete categorical distribution over *o_t_* with probability parameters resulting from transforming the distribution of *r_t_* by the observation function *g*. Formally,

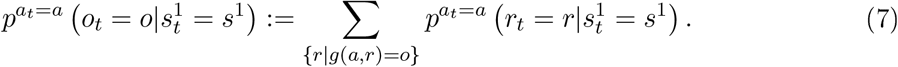

For the informative actions *a* ∈ {1, 3}, it thus follows that

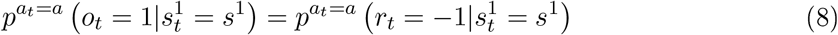

and

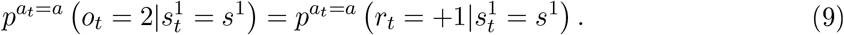

For the non-informative actions *a* ∈ {2, 4}, on the other hand, it follows that

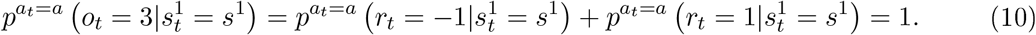

As an example, consider the case *s*^1^ = 1 (square is lucrative) and *a* = 1 (square on the informative side chosen). The agent allocates the same probabilities to observing either the image of the crossed-out moneybag or the image of the moneybag as to obtaining a reward of −1 or +1, respectively. Alternatively, consider the case *s*^1^ = 1 (square is lucrative) and *a* = 4 (triangle on the non-informative side chosen). In this case, the agent allocates a probability of 1 to observing the image of the question mark moneybag.

Based on the general agent structure encoded in the tuple *M*_Agent_, we next discuss our model space of interest, which comprises two control agents, denoted by C1 and C2, and three Bayesian agents, denoted by A1, A2, and A3.

##### Control agents Cl and C2

The control agents C1 and C2 rely on heuristic choice strategies. Because C1 and C2 do not represent a belief state, their action valence function is a function of action only,

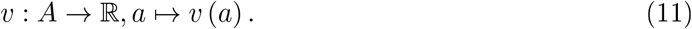

To realize an action on trial *t*, both agents use a probabilistic decision rule. Specifically, C1 and C2 directly translate the action valences into action and observation history-dependent choice probabilities.

##### Cl: A belief state-free random choice agent

Agent C1 may be considered a cognitive null model. It does not have an optimization aim based on which it could differentiate between actions, but merely allocates equal valences to all available actions 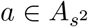,

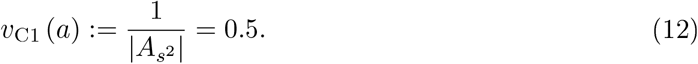

##### C2: A belief state-free win-stay-lose-switch agent

Agent C2 aims to maximize immediate rewards without relying on a belief state. To this end, C2 adopts a heuristic win-stay-lose-switch strategy (Robbins, 1952). Specifically, on each trial *t*, agent C2 determines its preferred choice based on previous reward signaling observations, but does not take action informativeness (i.e., shape laterality) into account. Formally, on trial *t* = 1, the action valence function of agent C2 is defined as

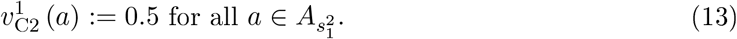

Subsequently, on trials *t* = 2, 3, …, *T* agent C2 allocates action valences according to

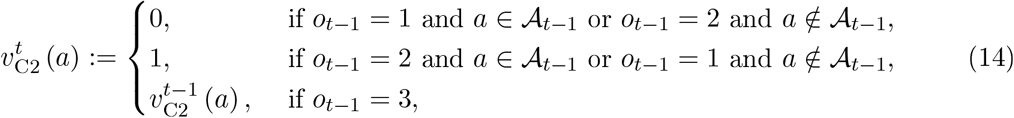

where 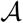 denotes the set of actions of choosing a given shape and thus 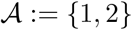 for the actions choose square and 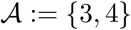 for the actions choose triangle.

Informally, agent C2 allocates equal initial action valences to all actions, because on trial *t* = 1 no previous observations are available and therefore the agent has no basis for differentiating between actions. Subsequently, on trials *t* = 2, 3, …, *T*, agent C2 allocates action valences depending on the observation on trial *t* − 1. Specifically, if on trial *t* − 1 the choice of a shape resulted in the observation *o* = 1, i.e., the image of the crossed-out moneybag, then agent C2 allocates an action valence of 0 to choosing the same shape and an action valence of 1 to choosing the other shape on trial *t*. In contrast, if on trial *t* − 1 the choice of a shape resulted in the observation *o* = 2, i.e., the image of the moneybag, then agent C2 allocates an action valence of 1 to choosing the same shape and an action valence of 0 to choosing the other shape on trial *t*. Crucially, if on trial *t* − 1 the choice of a shape resulted in the observation *o* = 3, i.e., the image of the question mark moneybag, the value of the returned reward is not signaled to the agent. In this case, agent C2 relies on its action valence allocation scheme from trial *t* − 1: The action valences that agent C2 allocates to choosing a given shape on trial *t* correspond to the valences the agent allocated to choosing that shape on trial *t* − 1.

##### Bayesian agents (Al, A2 and A3)

The Bayesian agents maintain a belief state which subserves their action choice. Specifically, the distributions 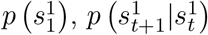 and 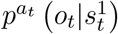 of *M*_Agent_ induce a trial-by-trial action-dependent joint probability distribution 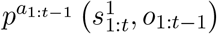. This allows for the recursive evaluation of the belief state 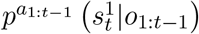 on trial *t* given the history of observations *o*_1:*t*−1_ and actions *a*_1:*t*−1_ by means of

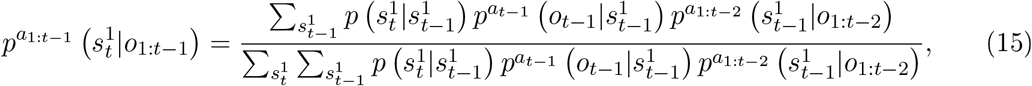

for *t* = 1, …, *T*, and where the prior belief state is given by 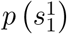 for *t* = 1. For the derivation of eq. (15), please see Section S.4: Belief state, posterior predictive distribution, and KL-divergence. Intuitively, the Bayesian agents update their belief state in a trial-by-trial fashion based on the observation made after choosing a shape on either side and by accounting for a reversal in the shapes’ lucrativeness. Note that on an implementational level, we represent the distributions 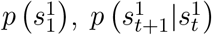 and 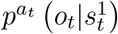 by stochastic matrices and evaluate the belief state using matrix multiplication as detailed in Section S.5: Belief state and posterior predictive distribution implementation.

Based on their belief state representation, the Bayesian agents then decide for an action based on a combination of an action valence function, which evaluates the desirability of a given action in the light of the agent’s current belief state, and a decision function, which selects the maximal desirable action as the action to issue. Specifically, the scalar representation of the belief state

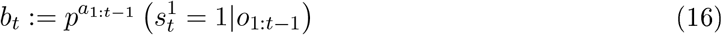

constitutes the basis for action evaluation by means of an action valence function

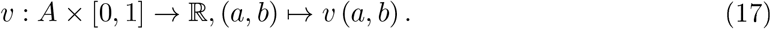

As detailed below, the exact forms of the valence function differ between agents A1, A2, and A3. However, to realize an action, all Bayesian agents pass the evaluated action valences on to a maximizing decision rule of the form

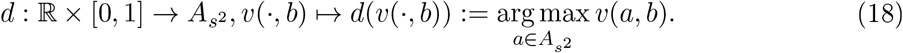

On every trial, the Bayesian agents thus choose the action with the highest valence.

##### Al: A belief state-based exploitative agent

Agent A1 uses its belief state to maximize the immediate reward gain. To this end, agent A1 uses an action valence function that allocates to action *a_t_* = *a* an action valence based on the action-dependent expected reward under the current belief state *b_t_* = *b* according to

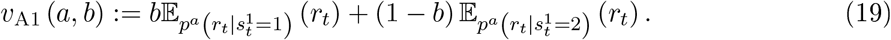

The upper and lower panels of Figure 3a visualize the A1 valences for actions *a* ∈ *A*_1_ (choose square on the informative side or triangle on the non-informative side) and *a* ∈ *A*_2_ (choose square on the non-informative side or triangle on the informative side), respectively, as functions of the belief state *b*. Note that the expected reward is

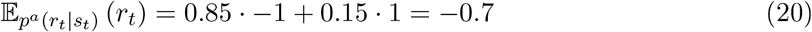

for choosing the detrimental shape and

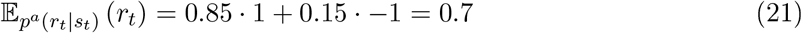

for choosing the lucrative shape. Consequently, the more certain A1 becomes that a given shape is lucrative (as *b* gets closer to 0 or 1 from 0.5) the higher the belief state-weighted expected reward for choosing that shape and, accordingly, the lower the belief state-weighted expected reward for choosing the other shape. As the belief state-weighted expected reward is irrespective of the side of the shape, in the case of both sets of available actions, agent A1 allocates valences without taking the actions’ informativeness into account.

**Figure 3.**
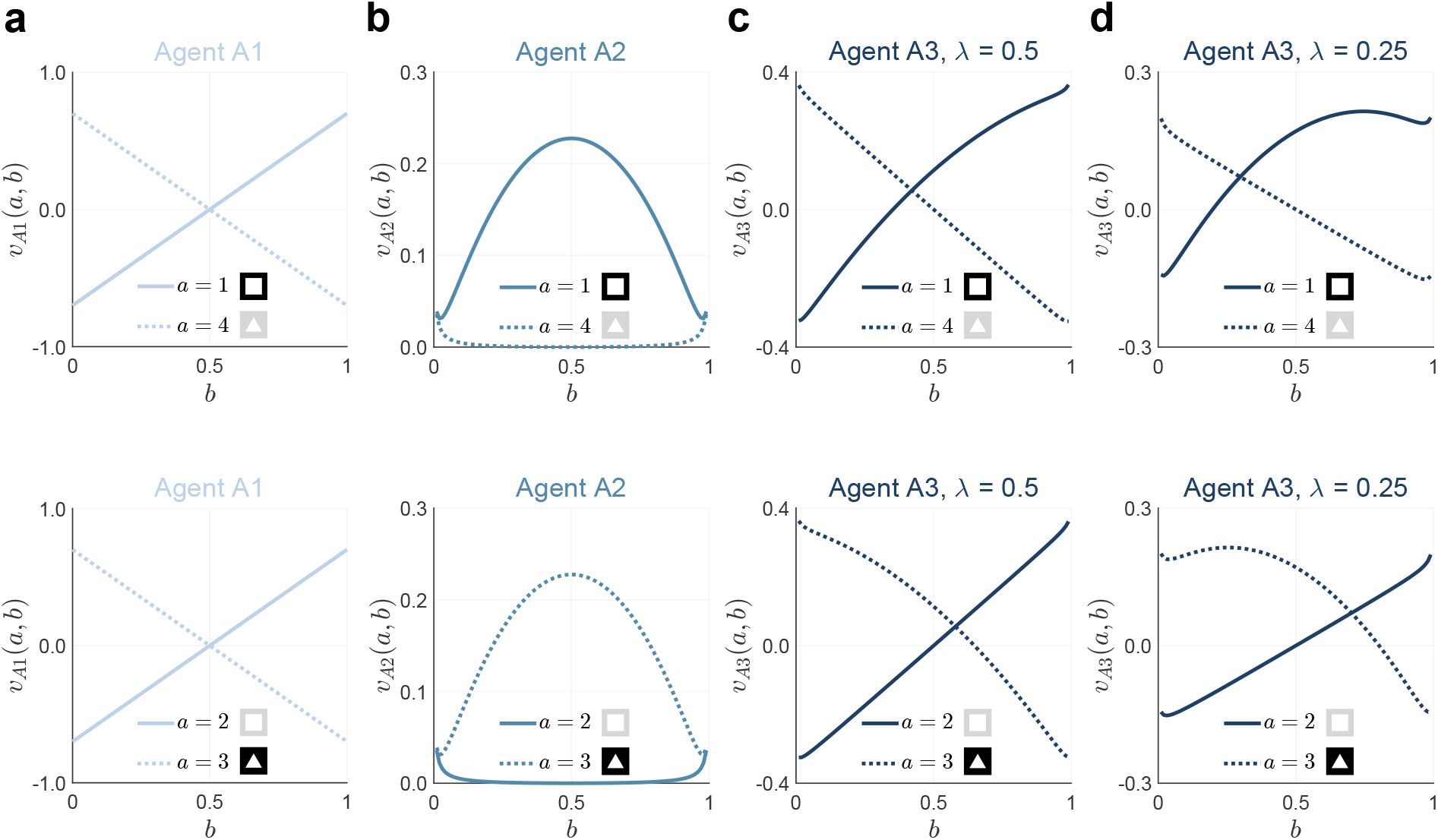
Bayesian agents’ action valence functions. **a** Action valence functions of agent A1 for the available action set *A*_1_ ≔ {1, 4} (upper panel) and the available action set *A*_2_ ≔ {2, 3} (lower panel). Agent A1 allocates action valences based on the belief state-weighted expected reward. As the expected rewards for choosing the lucrative or detrimental shape are constant, the more extreme the agent’s belief that a given shape is lucrative the higher the valence it allocates to choosing the corresponding shape and the lower the valence it allocates to choosing the other shape. The valences of A1 do not depend on the actions’ informativeness, which reverses between available action sets *A*_1_ and *A*_2_ and therefore the two panels are identical. **b** Action valence functions of agent A2 for the available action set *A*_1_ ≔ {1, 4} (upper panel) and the available action set *A*_2_ ≔ {2, 3} (lower panel). Agent A2 allocates action valences based on the expected Bayesian surprise, which is higher for the informative action than for the non-informative action, thus the graphs on the upper and lower panels flip. The higher the agent’s uncertainty about the lucrative shape, the larger the benefit of the informative action. **c, d** Action valence functions of agent A3 for the available action set *A*_1_ ≔ {1, 4} (upper panel) and the available action set *A*_2_ ≔ {2, 3} (lower panel) for *λ* = 0.5 (**c**) and *λ* = 0.25. Agent A3 allocates action valences based on the convex combination A1 and A2 action valences. The higher the value of *λ* the more the valences of A3 resemble the valences of A1 and correspondingly, the lower the value of *λ* the more the valences of A3 resemble the valences of A2. Note that the colors used for the graphical depiction of the agents’ action valence functions correspond to the agent model color scheme used for model recovery and model comparison in all Figures below. For implementational details, please see *figure_3.m*.

##### A2: A belief state-based explorative agent

Agent A2 explores its belief state to maximize the immediate information gain. To this end, on trial *t* agent A2 allocates a valence to each available action *a_t_* = *a* based on its expected Bayesian surprise

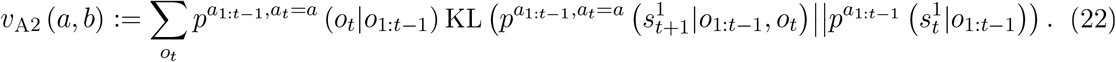

The first term in eq. (22),

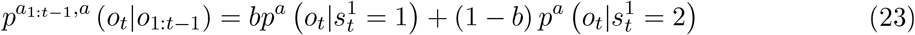

denotes the agent’s belief state-dependent posterior predictive distribution, which specifies the agent’s subjective uncertainty over the observation *o_t_* given action *a* on trial *t* and its history of observations *o*_1:*t*−1_ and actions *a*_1:*t*−1_. For a derivation of the right-hand side of eq. (23), please see Section S.4: Belief state, posterior predictive distribution, and KL-divergence and for implementational details regarding its evaluation, please see Section S.5: Belief state and posterior predictive distribution implementation. The second term in eq. (22),

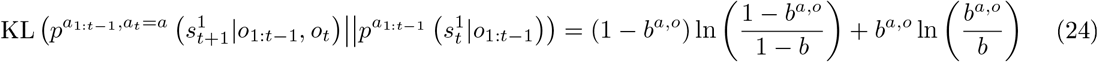

denotes the Kullback-Leibler (KL) divergence between the agent’s belief state on trial *t* and its virtual belief state on trial *t* + 1 given observation *o_t_* and action *a_t_* = *a* on trial *t*. Intuitively, this KL divergence quantifies the information gain afforded by choosing action *a_t_* = *a* and making observation *o_t_* on trial *t*. On the right-hand side of eq. (24), *b* denotes the agent’s belief state on trial *t* (cf. eq. (19)) and *b^a,o^* denotes the agent’s belief state *b_t_*_+1_ resulting from action *a_t_* = *a* and observation *o_t_* = *o*. For a derivation of the right-hand side of eq. (24), please see Section S.4: Belief state, posterior predictive distribution, and KL-divergence. In summary, *v*_A2_ (*a, b*) quantifies the agent’s weighted expected information gain afforded by choosing *a_t_* = *a* given its current belief state *b_t_* = *b*, where the expectation is formed with regards to the possible observations *o_t_* on trial *t* and the weighting is determined by the agent’s current estimate of observing a specific observation *o_t_* = *o*.

The upper and lower panels of Figure 3b visualize the A2 valences for actions *a* ∈ *A*_1_ (choose square on the informative side or triangle on the non-informative side) and *a* ∈ *A*_2_ (choose square on the non-informative side or triangle on the informative side), respectively, as functions of the belief state *b*. Choosing the shape on the non-informative side does not deliver reward information. Therefore, the expected Bayesian surprise-based A2 valence is always higher for the informative action, irrespective of the agent’s belief state. Yet, the difference between the informative and non-informative action valences depends on the belief state. Specifically, in contrast to agent A1, the more uncertain agent A2 becomes about the lucrative shape (as *b* gets closer to 0.5 from 1 or 0), the larger the difference between the valences and thus the stronger the agent’s preference for the informative action.

##### A3: A belief state-based explorative-exploitative hybrid agent

Agent A3 combines the choice strategies of agents A1 and A2 and uses its belief state to maximize the combination of immediate reward gain and information gain. Formally, on each trial *t* and for each available action 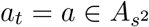, agent A3 evaluates its action valences based on the convex combination of the action valences of agents A1 and A2,

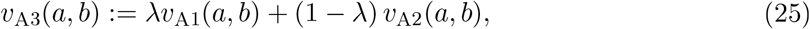

where *λ* ∈ [0, 1] is the weighting parameter. The upper and lower panels of Figures Figure 3c and Figure 3d visualize valences of agent A3 for actions *a* ∈ *A*_1_ (choose square on the informative side or triangle on the non-informative side) and *a* ∈ *A*_2_ (choose square on the non-informative side or triangle on the informative side), respectively, as functions of the belief state *b* for weight parameter values of *λ* = 0.5 and *λ* = 0.25, respectively. For *λ* = 1, the action valences of agent A3 correspond to the action valences of A1, while for *λ* = 0 the action valences of agent A3 correspond to the action valences of agent A2. For weighting parameter values *λ* ∈]0, 1[, the decision strategy of agent A3 results from a mixture of the strategies of agents A1 and A2: For non-extreme belief state values, i.e., *b* values close to 0.5, agent A3 allocates a higher valence to choosing the shape on the informative side, even if the agent allocates a lower probability to that shape being lucrative. This shows the contribution of agent A2’s choice strategy. For more extreme belief state values, i.e., *b* values close to 0 or 1, agent A3 allocates a higher valence to choosing the shape with the higher probability to be lucrative, even if the action is non-informative. This shows the contribution of agent A1’s choice strategy. Note, however, that a *λ* value of 0.5 should not be understood as the agent A3’s choice strategy resembling respective strategies of agents A1 of A2 to equal degree. The reason for this is that agent A3 applies a convex combination of the A1 and A2 action valences, which take values in different ranges (−0.7 to 0.7 for A1, 0 to 0.23 for A2). Therefore, while for *λ* = 0.5 the action valences of agent A3 primarily reflect the contribution of agent A1 (cf. Figure 3c), the contribution of the action valences of agent A2 becomes evident for *λ* = 0.25 (cf. Figure 3d).

#### Data analysis models

To evaluate the agent models in light of the participants’ data, we embedded the agent models in a statistical inference framework. In particular, for agent models C2, A1, A2, and A3 we formulated behavioral data analysis models by nesting the agent-specific action valence functions in a softmax operation (Reverdy & Leonard, 2015). To this end, we defined the probability of action *a* given the history of actions *a*_1:*t*−1_ and observations *o*_1:*t*−1_ as

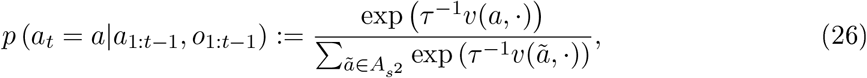

where for each agent, *v*(*a*, ·) corresponds to the agent-specific action valence function. Here, the parameter *τ* ∈ ℝ_>0_ encodes the level of post-decision noise: The lower the value of *τ*, the higher the probability that the action with the higher action valence is realized and thus the lower the post-decision noise. Notably, as agent C1 allocates equal action valences throughout, for any *τ* value the softmax operation returns a uniform probability distribution. Therefore, for this agent a softmax operation is not required, and we defined the respective data analysis model as

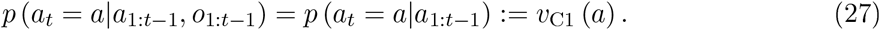

### Model estimation and comparison

To estimate the parameters of the data analysis models for agents C2, A1, A2, and A3, we used a maximum likelihood (ML) approach. Specifically, we assumed conditionally independently distributed actions and for each participant’s data defined the conditional log likelihood function as

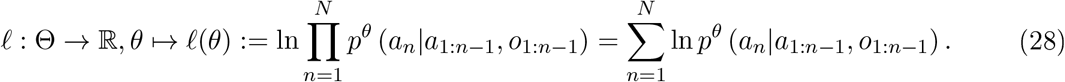

For agents C2, A1, and A2, the conditional log likelihood function is a function of the softmax operation parameter *τ* only. Thus, for these data analysis models *θ* ≔ *τ*. For agent A3, the conditional log likelihood function is a function of *τ* and the action valence weighting parameter *λ* ∈ [0, 1] (cf. eq. (25)). Thus, for this data analysis model *θ* ≔ (*τ, λ*). Note that in eq. (28) we index trials by *n* = 1, …, *N* rather than by *t* = 1, …, 2*T* to account for the fact that participant’s occasionally failed to respond, resulting in invalid choice trials. *n* hence indexes a participant’s *n*th valid choice trial and *N* denotes a participant’s total number of valid choice trials.

For every data analysis model, we estimated *θ* based on single participant data sets by minimizing the respective negative conditional log likelihood function using Matlab’s constrained nonlinear optimization function *fmincon.m* (Byrd et al., 1999, 2000; Waltz et al., 2006). The parameter constraints were set to *τ* ∈ [0.01, 2.5] and to *λ* ∈ [0, 1]. To mitigate the risk of identifying local maximizers rather than global maximizers of the conditional log likelihood function, the initial parameter estimate values were sampled from a continuous uniform distribution covering the constrained parameter space, the parameter estimation procedure was repeated ten times, and only the parameter estimates achieving the highest conditional log likelihood function value were regarded as ML estimates (Wilson & Collins, 2019). Note that the conditional log likelihood function of agent model C1 does not require any parameter optimization and can be evaluated directly.

To compare the models’ relative plausibilities given the participant’s data, we first evaluated all agent model- and participant-specific Bayesian Information Criterion scores according to (Schwarz et al., 1978)

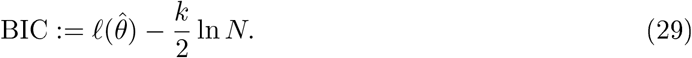

Here, 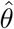 denotes the ML parameter estimate, *k* denotes the respective model’s number of to be estimated parameters, and *N* denotes the respective participant’s total number of valid choices. The BIC scores of all agents and participants were then subjected to a random-effects Bayesian model selection procedure as implemented in *spm_BMS.m* and distributed with the SPM toolbox for neuroimaging data analysis (www.fil.ion.ucl.ac.uk/spm/, Stephan et al. (2009); Rigoux et al. (2014)). For each model, *spm_BMS.m* returns a protected exceedance probability (PEP) which corresponds to the group-level probability that the particular model is more likely than the other models of the model space.

### Model and parameter recovery analyses

#### Model recovery analyses

To validate our agent-based behavioral modeling approach, we performed a number of model recovery simulations with the aim of assessing to which degree our analysis approach allows for reliably arbitrating between the models in our model space. To this end, we first generated synthetic behavioral data using each of our agent-based data analysis models C1, C2, A1, A2, and A3. The synthetic behavioral data sets comprised agent-specific actions on 160 trials under the experimentally employed state sequence (cf. Section S.3) and for agent action-dependent observations *o*_1:2*T*_ and rewards *r*_1:2*T*_. For agent C1, the synthetic data were generated without further specifications. For agents C2, A1, A2, and A3, the synthetic data were generated using softmax operation parameter values between *τ* = 0.05 and *τ* = 2.5 in steps of Δ*τ* = 0.05. In addition, for agent A3, the synthetic data were generated with parameter values *λ* ∈ {0.1, 0.25, 0.3, 0.5, 0.7, 0.9}. For each data generating model and for each parameter value, we generated 24 data sets (corresponding to the experimental sample size), evaluated these data sets using all models in our model space by means of the ML estimation and BIC model comparison approach discussed above, and repeated this procedure 10 times. Finally, as our primary outcome measure, we evaluated the average protected exceedance probabilities across the 10 data generation and data analysis repeats.

Figure 4 summarizes the results of the model recovery analyses. For each data generating model, the corresponding panel depicts the protected exceedance probabilities (PEPs) of each data evaluation model in the model space. As shown in the leftmost panel of Figure 4a, for data generated based on the cognitive null agent C1, the PEP is maximal for the data analysis model based on agent C1, thus indicating that agent model C1 can be reliably recovered. For the data generating models based on agents C2, A1, A2, and A3, the PEPs are shown as functions of the post-decision noise parameter *τ* used for data generation. For data generated based on agents C2 and A1, the PEP is maximal for data analysis models based on agents C2 and A1, respectively, for all values of *τ*. This indicates that agent models C2 and A1 can reliably be recovered for both low and high levels of post-decision noise. For data generated based on agent A2, the PEP is maximal for the data analysis model based on agent A2 for *τ* values up to 0.35, while the PEP is maximal for the data analysis model based on C1, if *τ* > 0.35. This indicates that agent model A2 can be reliably recovered for low, but not for high, levels of post-decision noise. For data generated based on agent A3 and with action valence weighting parameter set to *λ* = 0.25, the PEP is maximal for the data analysis model based on A3 up to a value of *τ* = 0.25. For larger values of *τ*, the PEP of the data analysis model based on agent A1 and eventually of the PEP of the data analysis model based on agent C1 exceed that of agent A3. This indicates that for *λ* = 0.25, the data analysis model based on agent A3 can be recovered reliably only for low levels of post-decision noise. With increasing noise, the data is better accounted for by the less complex model based on agent A1 and eventually by the cognitive null model C1.

**Figure 4.**
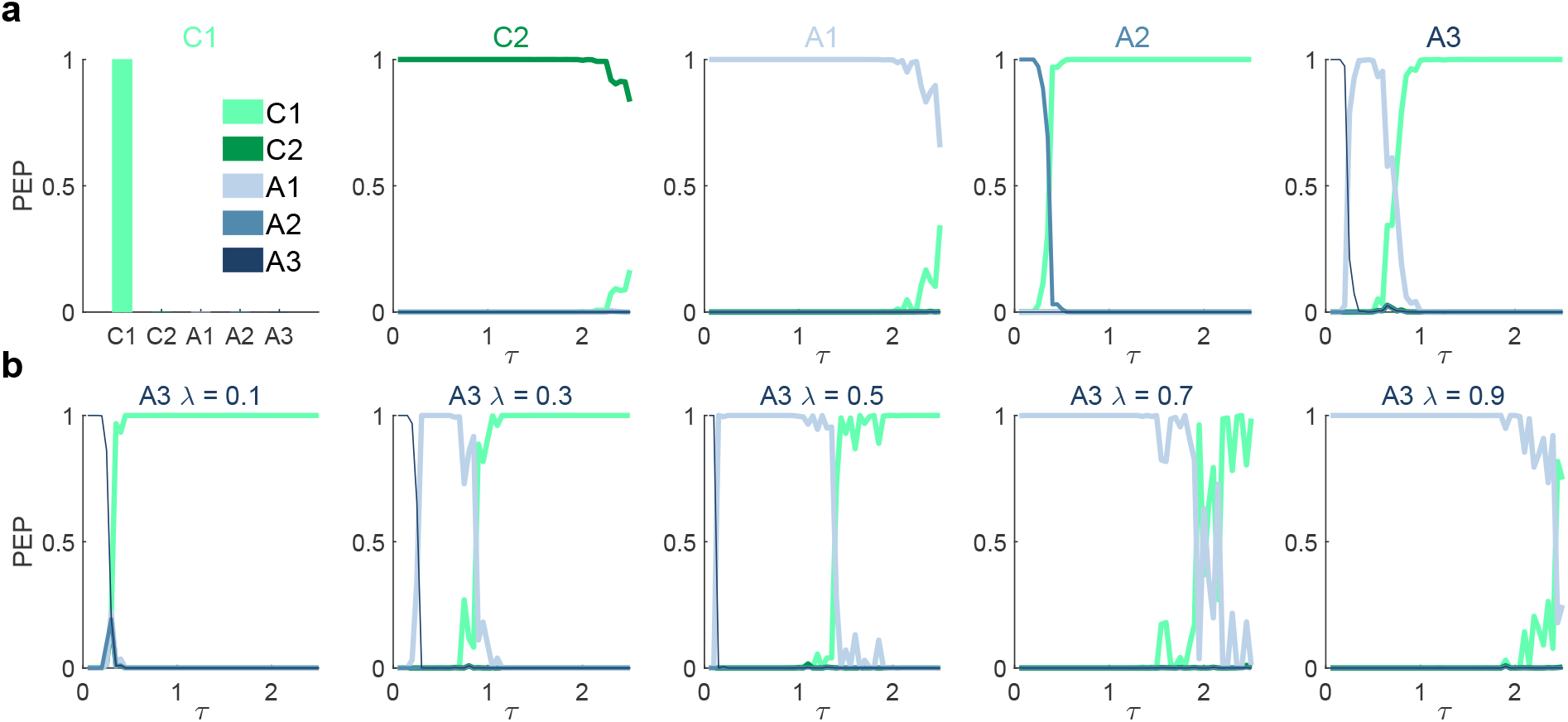
Model recovery results. **a** Model recovery results. Each subplot pertains to a specific data generating model (C1, C2, A1, A2, and A3) and shows the protected exceedance probabilities (PEPs) of all data analysis models in the model space evaluated in light of data sets sampled from the respective data generating model. For data generated based on the cognitive null model C1, the PEP was maximal for C1. For data generated based on agents C2, A1, A2, and A3, the PEPs depend on the value of the post-decision noise parameter *τ*. Agents C2 and A1 are recoverable up to high levels of post-decision noise. Agents A2 and A3 (for *λ* = 0.25) are recoverable only for low levels of post-decision noise. **b** Model recovery results for agent A3 for different values of the action valence weighting parameter *λ*. Agent A3 is recoverable up to medium values of *λ* and for low levels of post-decision noise. For implementational details, please see *figure_4.m*.

For data generated based on agent model A3, model recovery performance depends not only on the post-decision noise parameter *τ*, but also on the action choice weighting parameter *λ*. As shown in Figure 4b, for *λ* = 0.1, *λ* = 0.3, and *λ* = 0.5 and for small values of *τ*, the PEP is maximal for the data analysis model based on agent A3. For *λ* = 0.1 and increasing post-decision noise, first the data analysis based on agent A2 and next the cognitive null model C1 prevail (left-most panel). For *λ* values of *λ* = 0.7 and *λ* = 0.9, the PEP profiles shift towards a prevailment of the model based on agent A1, in line with the increasing similarity between the action valences of agents A3 and agent A1 for increasing values of *λ* (cf. Figure 3c-d). More precisely, for *λ* = 0.7 and *λ* = 0.9 the PEP is maximal for model A1 up to values of *τ* = 1.9 and *τ* = 2.4, respectively. For even larger values of *τ*, the PEP of the cognitive null model C1 prevails. Together, in line with the meaning of the action valence weighting parameter of agent A3, these results indicate that agent model A3 is reliably recoverable for low to medium values of *λ* in the presence of low levels of post-decision noise. For very low values of *λ* and low levels of *τ*, model A3 cannot be distinguished from model A2, while for high levels of *λ*, it cannot be distinguished from model A1. Finally, for high levels of post-decision noise, the cognitive null model C1 provides the most parsimonious data explanation for data generated by agent model A3.

#### Parameter recovery analyses

Additionally, to assess to which degree we can reliably estimate the data analysis model parameters, we performed a number of parameter recovery analyses. Specifically, for a given data generating model and model parameter setting, we evaluated the average ML parameter estimates and their associated standard errors of the data generating model across the 24 synthetic data sets and the 10 data generation and data analysis repeats.

Figure 5 summarizes the parameter recovery analyses results. Each panel of Figure 5 depicts model-specific ML parameter estimates as functions of the post-decision noise parameter *τ*. Specifically, Figure 5a depicts the parameter estimate 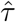 as a function of the respective data generating parameter values *τ* under the data generating and data analysis models C2, A1, A2, and A3 with *λ* = 0.25. In general, the parameter estimates are consistent with their respective data generating parameters for small values of *τ*. For larger values of *τ*, the true parameter values are first over- and then underestimated. As shown in Figure Figure 5a, this bias is subtle for agents C2 and A1 and only appears for relatively large *τ* values above *τ* = 1.5. For agents A2 and A3 the bias is more pronounced and affects medium to large values above *τ* = 0.5. These results are consistent with the model recovery results: For large post-decision noise, data generated by any model starts to resemble data generated by a purely random choice strategy and thus the parameter estimate for *τ* reaches an asymptote.

**Figure 5.**
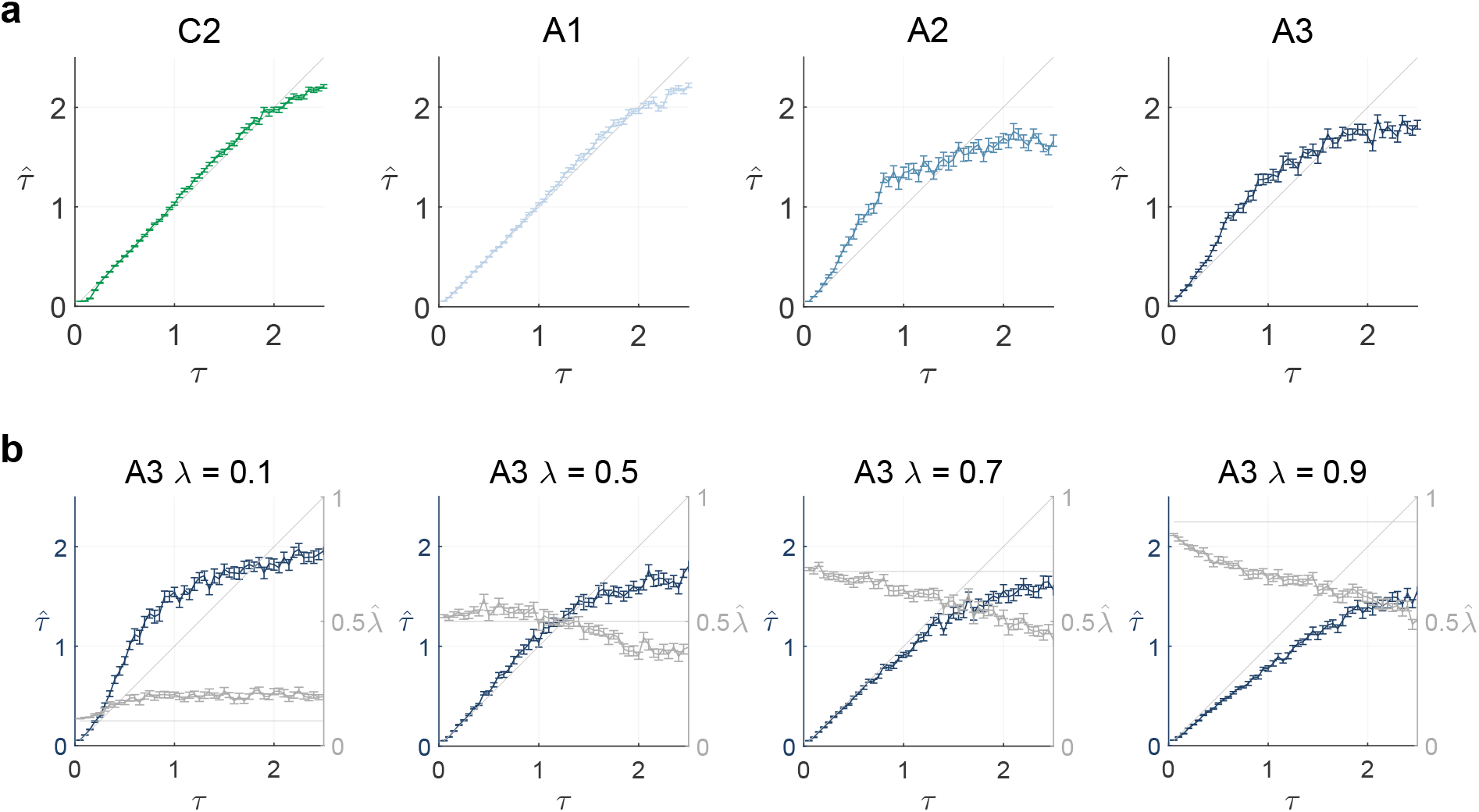
Parameter recovery results. **a** Parameter recovery results for all agents with free parameters. For every agent, the corresponding panel shows the ML parameter estimates as functions of the post-decision noise parameter *τ* used for data generation. The post-decision noise parameter of agents C2 and A1 is recoverable from small to medium values. The post-decision noise parameter of agents A2 and A3 with *λ* = 0.25 is recoverable for small values. **b** Parameter recovery results for agent A3 with different *λ* values. The parameter *τ* is recoverable for *τ* values up to *τ* = 1, upon which it exhibits a downward bias. The parameter *λ* is recoverable for small *τ* values, except for a slight downward bias for *λ* = 0.9. For medium to large values of the post-decision noise parameter, 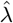 is downward biased. For implementational details, please see *figure_5.m*.

The panels of Figure 5b show the parameter estimates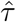 and 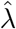 of agent model A3 as a function of *τ* for *λ* = 0.1, *λ* = 0.5, *λ* = 0.7, and *λ* = 0.9, respectively. As above, the parameter estimates 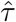 show a pattern of consistent estimation for values up to *τ* = 1, upon which they exhibit a downward bias. The parameter *λ* is reliably recovered for small values of *τ*, except for a slight downward bias for *λ* = 0.9. However, for medium to large *τ* values, 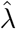 is more strongly downward biased. These findings are consistent with the model recovery analyses results: First, the deflation effect for *λ* = 0.9 and small values of *τ* shows that for large values of *λ*, data generated by agent A3 is virtually indistinguishable from data generated by agent A1 and thus 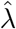 reaches an asymptote. Second, the bias of 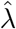 for medium to large values of *τ* reiterates that with increasing post-decision noise, data generated by agent A3 increasingly resembles data generated by a random choice strategy, such that *λ* cannot be reliably identified.

### Model comparison results

Upon validating our modeling initiative, we evaluated and compared our agent-based model set in light of the participants’ data. For 18 of the 24 participant data sets, the BIC score was maximal for the model based on agent A3. Accordingly, as shown in Figure 6a, the group cumulative BIC score was maximal for model A3 and the group-level PEP of model A3 was larger than 0.99. The ML parameter estimates 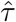 and 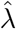 for model A3 varied moderately across participants. Specifically, 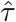 ranged from 0.035 to 0.377 with an average of 0.124 ± 0.014 and 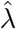 ranged from 0.044 to 0.622 with an average of 0.274 ± 0.027. Notably, these are parameter ranges, in which A3 is well recoverable and identifiable. Taken together, these findings indicate that agent model A3 provides the most plausible explanation of the participant’s choices among the set of behavioral models assessed and that the most frequently applied choice strategy among the group of participants resembled that of agent A3 most.

**Figure 6.**
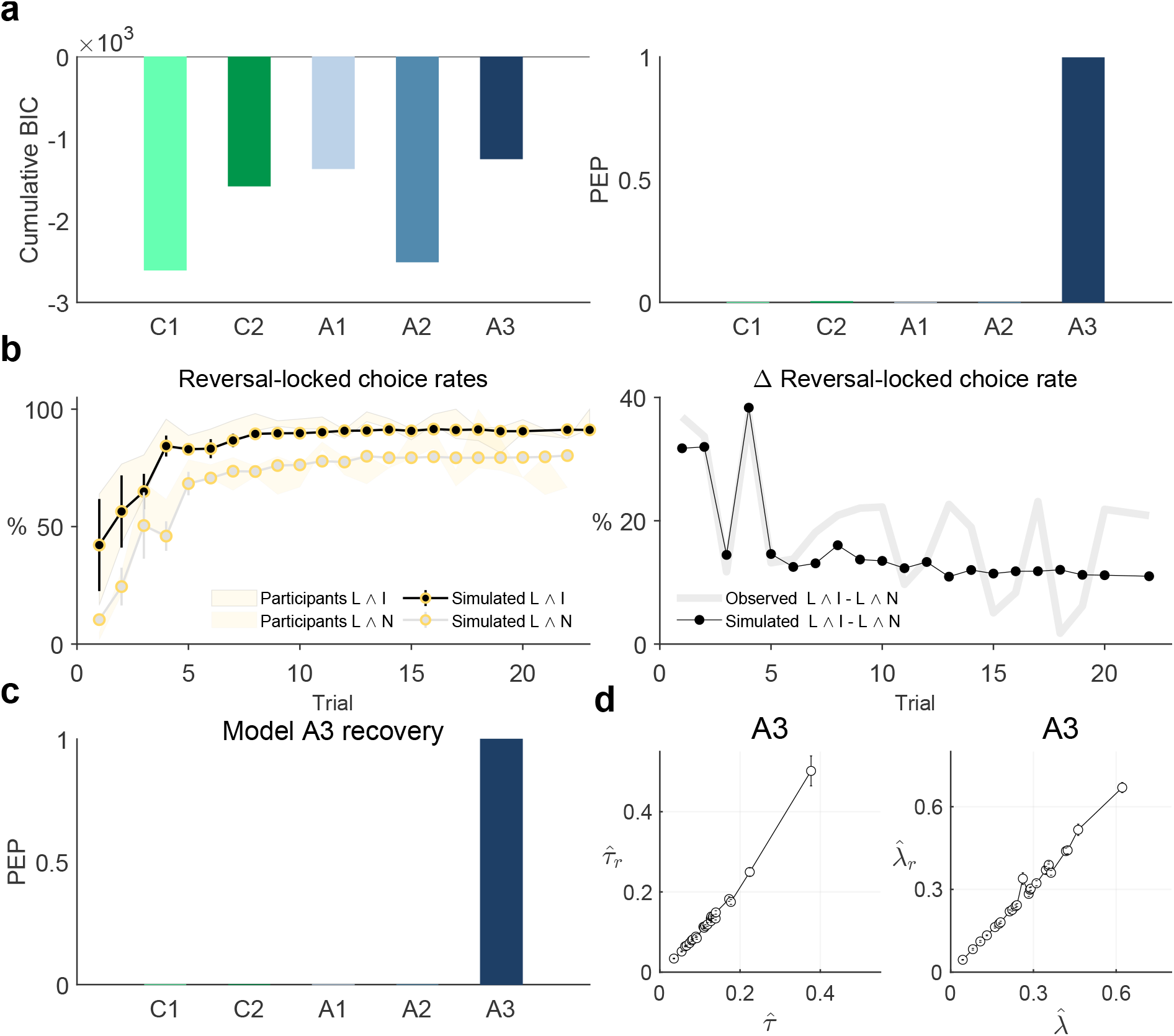
Computational modeling results. **a** Model comparison results. Both the group cumulative Bayesian information criterion scores (cumulative BIC, left panel) and the protected exceedance probabilities (right panel) were maximal for the agent model A3, indicating that this model explained participants’ choice data the best. **b** Model A3 behavioral validation. Average reversal-locked group trial-by-trial L ∧ I action and L ∧ N action choice rates (left panel) and their difference (right panel) based on synthetic data sets generated using agent A3 and the participants’ parameter estimates. The patterns closely resemble those observed in the participants’ data (cf. Figures 3c, 3d), re-visualized here for convenience. **c** Model A3 recovery based on the participants’ data parameter estimates. The protected exceedance probability based on re-analysis of simulated data from agent model A3 with parameters set to the participants’ parameter estimates is maximal for agent model A3. **d** Parameter recovery results based on data generated with agent model A3 and the participants’ data-based parameter estimates. The panels depict the simulation-averaged participant-specific recovered parameter estimates 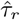 and 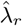 and their associated SEMs over simulations as a function of the participant-specific parameter estimates 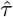 and 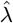. Both the participants’ post-decision noise parameter estimates 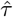 as well as the participants’ weighting parameter estimates 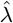 are recoverable. For implementational details, please see *figure_6.m*.

#### Post-hoc model validation

Upon identifying agent A3 as the most plausible model to explain our observed experimental data based on the formal criteria discussed above, we further investigated the behavioral face validity and the participant parameter estimate-based identifiability of this model. To this end, we first generated 24 synthetic data sets based on agent A3 using the 24 participant-specific ML parameter estimates. As in the model recovery analysis, the synthetic data conformed to the agent’s actions on 160 trials, with trial sequence identical to the participants’ trial sequence and with agent action-dependent observations. To assess the model’s behavioral face validity, we then subjected these 24 data sets to the descriptive analyses described above and computed the same action choice rates as for the participants’ data. To minimize random variation, we repeated this process 100 times and for each action choice rate evaluated the average over the 100 simulations. Similarly, we subjected the simulated data sets to our model estimation and comparison procedures and evaluated the average model and parameter recoverability performance as discussed above. We next evaluated the same summary and trial-by-trial choice rates as for the participants’ data. Consistent with the empirically observed results, most synthetic action choices were lucrative and informative (L ∧ I, 84.98% ± 1.29), followed by significantly fewer lucrative and non-informative synthetic actions (L ∧ N66.35%±1.96, two-sided paired sample t-test, *t*(23) = 11.59, *p* < 0.001). Furthermore, as shown in Figure 6b the reward reversal-locked trial-by-trial dynamics of the synthetic actions exhibited a very similar pattern to that of the participants (cf. Figure 2c,d). Specifically, while both the average L ∧ I as well as the average L ∧ N action choice rates increased between two reversals (left panel), their difference decreased moderately between the first trial after and the last trial before a reward rate reversal (right panel). Taken together, these results support the behavioral validity of the A3 model. At last, we conducted model and parameter recovery analyses for model A3 based on the parameter estimates for the 24 participant data sets. As already implied by the results of the full parameter space recovery analyses reported above, these analyses confirmed that also for the range of empirically estimated parameter values, both the model and its parameters are reliably recoverable (Figure 6c and Figure 6d).

## Discussion

In this work, we addressed the question of how humans make sequential decisions if all available actions bear economic consequences, but only some also deliver choice-relevant information. By collecting human choice data on an information-selective symmetric reversal bandit task, we demonstrated that in such situations, humans strive for a balance of exploratory and exploitative choice strategies. To arrive at this conclusion, we applied a set of descriptive and agent-based computational modeling analyses, including model evaluation based on formal as well as informal criteria. Given our model set, the behaviorally most plausible strategy was captured by a Bayesian agent that assessed the desirability of an action by applying a convex combination of its expected Bayesian surprise and its expected reward. A series of recovery analyses validated our modeling initiative and supports the robustness and reliability of our results. In sum, the key contributions of this work are threefold: We designed an information-selective symmetric reversal bandit task, we formulated, implemented, and validated a set of agent-based behavioral models that interact with this task, and we provide empirical evidence for human uncertainty-guided exploration-exploitation choice strategies. In the following, we discuss each of these contributions in turn.

As the first of our contributions, we introduced an information-selective symmetric reversal bandit task suitable to model a class of sequential decision-making problems, in which information about the conferred reward is not available for every action. In the following, we briefly discuss the key aspects of our task with respect to other tasks in the literature. As alluded to in the Introduction, previous research primarily employed either classical bandit paradigms or pure-exploration paradigms to study sequential decision making under uncertainty (Bubeck et al., 2009; Hertwig & Erev, 2009; Ostwald et al., 2015; Sutton & Barto, 2018; Wulff et al., 2018). We consider these paradigms in turn.

In the classical bandit paradigm, the deciding agent chooses between a set of ‘arms’ (Berry & Fristedt, 1985). Similar to our task, each arm confers reward according to its associated reward distribution, and, in contrast to our task, each arm confers also information about its expected reward value. A drawback of this design is that the co-occurrence of reward and information evokes a confound between an action’s value estimate and the associated uncertainty: as people tend to favor the action with the higher value estimate, for that action the associated uncertainty becomes smaller - simply because for that action more reward observations are made. This makes it difficult to uncover uncertainty-guided exploration in the classical bandit paradigm (Dezza et al., 2017; Gershman, 2018; Wilson et al., 2014). The task employed in the current study circumvents this problem by adopting a symmetrical reward structure of the actions: the probability of the positive reward for the lucrative action is identical to the probability of the negative reward for the detrimental action. Likewise, the probability of the negative reward for the lucrative action is identical to the probability of the positive reward for the detrimental action. In this manner, each reward observation following the informative action confers the same amount of information about the expected reward value of both the lucrative and detrimental action. Furthermore, as in each trial information is randomly coupled with either the lucrative or the detrimental action, our task arguably evokes a more explicit exploration-exploitation dilemma than the classical bandit paradigm, in particular on trials on which participants face decisions between potentially lucrative, but non-informative and detrimental, but informative actions.

The pure-exploration paradigm models sequential decision-making problems in which an action either confers information or reward. In the classical variant of this paradigm, an action that confers reward automatically terminates the task. In an extended variant, referred to as the ‘observe-or-bet task’ (Blanchard & Gershman, 2018; Navarro et al., 2016; Tversky & Edwards, 1966), the deciding agent can freely switch between an ‘observe action’ that confers information and ‘bet actions’ that confer reward. Specifically, ‘observe actions’ return information about the expected reward of the bet actions, but do not return reward. ‘Bet actions’, on the other hand, return rewards according to their associated reward distributions, but no information. Similar to the bet actions, in our task one of the available actions confers only reward, but no information. However, in our task, the other available action does not only confer information, as the ‘observe action’ does, but it also confers reward. Therefore, while exploration and exploitation are temporally separated in the observer-or-bet task, they have to be balanced simultaneously in our task.

In summary, the task proposed in the current study complements the experimental arsenal for studying human exploration-exploitation behavior in the following sense: In contrast to the classical bandit paradigm, the exploration-exploitation dilemma on each trial is *explicit*, i.e., participants need to actively decide whether to gather reward and/or information, and do not obtain information as a by-product of reward-maximizing decisions as in the classical bandit task. Further, in contrast to the pure-exploration and observe-or-bet task, exploration and exploitation phases occur *simultaneously* and not separated in time. Finally, by introducing a constant probability for the reward rate reversal of the choice options, the task does not only evoke an explicit and simultaneous exploration-exploitation dilemma, but also one that is *ongoing*. This aspect is in line with recent studies using classical bandit or observe-or-bet tasks that also adopted non-stationary reward structures (Bartolo & Averbeck, 2020; Blanchard & Gershman, 2018; Chakroun et al., 2020; Navarro et al., 2016; Hampton et al., 2006; Speekenbrink & Konstantinidis, 2015) and emulates the ever-changing reward structure of real environments.

Our second contribution is the formulation and validation of a set of agent-based behavioral models that can interact with the information-selective symmetric reversal bandit task. Specifically, our model space comprises belief state-based agents that formalize subjective uncertainty-based exploitative, explorative and hybrid explorative-exploitative strategies, as well as belief state-free agents that formalize random choice and heuristic win-stay-lose-switch strategies. The belief state-based agents implement recursive belief-state updates to infer the not directly observable structure of the task environment, i.e., the most likely currently lucrative action. While this form of optimal Bayesian learning may seem to be a strong assumption, it has been shown to approximate human learning reasonably well in comparable switching state tasks, such as the two-armed reversal bandit task, non-stationary versions of the observe-or-bet task, and the multi-armed bandit task (Hampton et al., 2006; Blanchard & Gershman, 2018; Navarro et al., 2016; Daw et al., 2006; Speekenbrink & Konstantinidis, 2015). Moreover, by also including belief state-free agents in our model space, we also accounted for simple strategies that do not require Bayesian update. Of these, the win-stay-lose-switch agent adopts a well established effective strategy to approach similar bandit problems (Robbins, 1952).

The three belief state-based agents implement their respective choice strategies by following different optimization aims. In particular, the belief state-based exploitative agent A1 seeks to maximize the belief state-weighted expected reward. The belief state-based explorative agent A2 seeks to maximize the expected Bayesian surprise. The belief state-based hybrid explorative-exploitative agent A3 seeks to maximize the convex combination of these two quantities. Belief state-weighted expected reward is a natural quantity to formally capture immediate reward gain and thus myopic exploitation (Sutton & Barto, 2018). Expected Bayesian surprise is one of many quantities that have been proposed to formally capture immediate information gain and thus myopic exploration (Schwartenbeck et al., 2019). As alluded to in the Introduction, we here opted for Bayesian surprise due to its putative representation in the human neurocognitive system (Itti & Baldi, 2009; Ostwald et al., 2012; Gijsen et al., 2020).

Importantly, we would like to note that our definition of exploration here pertains to a form of exploration that is generally referred to as ‘directed exploration’ (Gershman, 2018, 2019; Wilson et al., 2014). This term is used to distinguish information gain maximizing exploration from ‘random exploration’. Random exploration is a form of exploration that achieves information gain by implementing a stochastic strategy, i.e. making stochastic choices based on the actions’ reward value estimate. While there are more principled ways such as Thompson sampling (Thompson, 1933), random exploration is commonly accounted for by the softmax operation (Reverdy & Leonard, 2015). Notably, we here do not interpret the softmax operation as random exploration. Instead, we employ the softmax operation to embed the agents in a statistical framework that accounts for post-decision noise. This way, we clearly separate the (deterministic) choice strategies implemented by the agents and the statistical agent-based behavioral data analysis models. In future work, we aim to broaden our model space by considering agents that adopt random exploration. Crucially, this necessitates a probabilistic filtering framework that allows to partition the variability of choice data into components relating to an agent’s stochastic exploration strategy and to post-decision noise (c.f. Ostwald et al., 2014).

As a third contribution, we provided evidence for human uncertainty-guided exploration-exploitation in the information-selective symmetric reversal bandit task: As uncertainty decreased, participants’ choices were less influenced by the prospect of information gain and more influenced by the prospect of reward gain. This finding is consistent with the behavior in the observer-or-bet task. In the first empirical study using the observer-or-bet task, (Tversky & Edwards, 1966) found that participants explored more, i.e., chose the observe action more frequently, if they (falsely) believed that the underlying environment was dynamic, i.e., the lucrative and detrimental bet actions reversed over time. While (Tversky & Edwards, 1966) did not relate this result to the notion that dynamic environments promote uncertainty, which, in turn, promotes exploration, in a recent study, (Navarro et al., 2016) formally tested this hypothesis. By modeling participants’ choices in both static and dynamic versions of the observer-or-bet task, they demonstrated that switches between the exploratory observe action and the exploitative bet actions were mediated by uncertainty. Our result is also in line with recent findings from studies employing multi-armed bandit tasks. Work by several authors showed that when controlling for the value estimate-uncertainty confound, behavior in static two-armed bandit tasks reflects an uncertainty-dependent combination of exploration and exploitation (Dezza et al., 2017; Gershman, 2018, 2019; Wilson et al., 2014). Notably, however, consistent with the notion that the value estimate-uncertainty confound has the potential to mask directed exploration, findings from earlier studies not accounting for this confounding effect are less conclusive. For example, (Zhang & Angela, 2013) also found evidence for a belief state-based explorative-exploitative strategy in static four-armed bandit tasks. In contrast, (Daw et al., 2006) did not find evidence for such a strategy in analyzing choices in a dynamic four-armed bandit task with changing action values. While the finding from (Daw et al., 2006) is contrary to our finding, acknowledging that value estimate and uncertainty are not confounded in our task, these two findings can be reconciled.

At last, some limitations of our study along with some suggestions for future research must be acknowledged. First, as for any scientific study, inferences can only be made with respect to the model space, which *per se* is incomplete. Thus, although we provide evidence for a strategy resembling a belief state-based agent seeking to maximize the convex combination of its expected Bayesian surprise and its expected reward, it is very well possible that an agent not included in our model space can better account for the behavioral data. Based on our experimental documentation and the open availability of the data, future research may broaden the model space and could also consider, for example, probabilistic variants of the win-stay-lose-switch agents (Worthy et al., 2012) or agents with non-adaptive learning rates (Wiering & Otterlo, 2012; Rescorla & Wagner, 1972), besides the already mentioned random exploration agents in a probabilistic filtering framework (Ostwald et al., 2014). Second, on a related note, we here do not derive an optimal agent in the sense of partially observable Markov decision process theory (Bertsekas, 2000; Puterman, 2005; Bäuerle & Rieder, 2011). A model space that comprises an agent implementing an optimal choice strategy would afford the analysis of the participants’ behavior from a normative perspective and is thus an interesting endeavor for future research. To facilitate such an undertaking, we have provided a detailed documentation of the task and agents developed thus far. Third, given the modest sample size of our study, the behavioral data reported here are best capitalized on in an exploratory fashion. Together with the implemented open research measures (Ritchie, 2020), we believe to have laid the foundations for reproducing our study and build upon our work.

### Conclusion

In conclusion, we here introduced a new behavioral task that models a subset of real-life sequential decision-making problems that has thus far received relatively little attention in the computational modeling literature: problems, in which information about the conferred reward is not available for every action. Importantly, this task allows to investigate a pronounced form of simultaneous exploration and exploitation processes without introducing a value estimation-uncertainty confound. By analyzing participants’ choices on this task using descriptive methods and agent-based behavioral models, we provide evidence for an uncertainty-guided balance between exploration and exploitation in human sequential decision making under uncertainty.

## Acknowledgements

We are grateful to Russell Tobe1 Melissa Breland1 Tina Bermudez1 and Alexis Lieval for help with participant recruitment1 Anna MacKay-Brandt for help with participant enrollment1 and Caixia Hu and Raj Sangoi for help with data acquisition. We thank Hauke Heekeren for support throughout this work.

## Declarations

### Ethics approval

The study was conducted in line with the human participant guidelines of the 1964 Declaration of Helsinki and was approved by the Institutional Review Board of the Nathan Kline Institute for Psychiatric Research (Orangeburg1 NY1 US).

### Consent to participate and consent for publication

Informed consent for participation and publication was obtained from all individual participants included in the study.

### Availability of data and material

The behavioral datasets generated and analysed during the current study are available from the Open Science Framework at https://osf.io/vdmah/.

### Code availability

All custom Python and Matlab R2020a (The MathWorks1 Natick1 USA) code implementing the experimental paradigm1 simulations1 and analyses is available from the Open Science Framework at https://osf.io/vdmah/.

### Authors’ contributions

L.H.: conceptualization, project administration, data curation, investigation, methodology, formal analysis, software, validation, visualization, writing - original draft, writing - review & editing; S.C.: project administration, data curation, investigation, experimental software, funding acquisition, resources, supervision, writing - review; M.M.: conceptualization, funding acquisition, resources, supervision, project administration, writing - review; S.R.: project administration, data curation, investigation, writing - review; P.S.: conceptualization, writing - review; D.O.: conceptualization, formal analysis, methodology, software, visualization, project administration, resources, validation, funding acquisition, supervision, writing - original draft, writing - review & editing.

## Supplementary material

### S.1. Sample characteristics

To characterize the group of participants, we measured symptoms of attention deficit hyperactivity disorder (ADHD), anxiety, depression and impulsivity. To this end, we used the questionnaires Conners Adult ADHD Rating Scale - Self Report, Short Version (CAARS-S:S; Conners et al. (1999)), State and Trait Anxiety Inventory (STAI; Spielberger et al. (1983)), Beck Depression Inventory II (BDI-II; Beck et al. (1996)) and UPPS-P Impulsive behavior Scale (Lynam et al., 2006), respectively. As shown in Table S.1, the sample varied only moderately with respect to these symptoms. For example, on CAARS-S:S, our main questionnaire of interest, participants scored within ±2 standard deviations of the mean of their age- and gender-matched norm groups of the general population. We therefore concluded that the sample represents the healthy population and did not relate individual variability in terms of ADHD or other clinical symptoms to behavioral strategies. We also report the IQ score, which was obtained by administering the Wechsler Abbreviated Scale of Intelligence (WASI-II; Wechsler (1999)) at the time of the Nathan Kline Institute Rockland Sample study (Nooner et al., 2012).

**Table S.1.** Sample characteristics.

| Measurement | Range | Median | Mean ± SD |
| --- | --- | --- | --- |
| Age (years) | 18 - 35 | 23.5 | 24.5 ± 5.53 |
| WASI-II (total score) | 84 - 122 | 101.5 | 102.38 ± 9.14 |
| CAARS-S:S (total T-score) | 32 - 65 | 48 | 47.63 ± 9.16 |
| BDI-II (total) | 0 - 20 | 4 | 6.67 ± 6.3 |
| STAI STATE (total T-score) | 34 - 63 | 43 | 45.6 ± 8.52 |
| STAI TRAIT (total T-score) | 34 - 71 | 49.5 | 49.92 ± 10.29 |
| UPPS-P (total) | 73 - 190 | 118.5 | 124.04 ± 25.66 |

### S.2. Participant instructions

Participants were provided with the following task instructions:

Welcome to the main part of today s experiment! In the following we will introduce to you the decision making task that you will complete in the scanner. Please read the instructions carefully. If you have any questions, feel free to ask at any time. Once you read the instructions, you will complete a test run with the task to make sure you feel comfortable with it before going in the scanner. On every trial we will present to you two objects, an orange square and a blue triangle on either side of a black and grey screen and ask you to choose between them. One of these objects is profitable, meaning that it is going to give you a win most of the time, while the other object in not profitable meaning that it is going to give you a loss most of the time. Once you choose one of the objects, the outcome (win: +1 or loss: −1) will be registered to your account. You will have 2.5 seconds to indicate your choice. If you do not respond within this time window, the message Too slow will appear on the screen and you automatically lose 1 point. Here you see an example for a trial (Supplementary Figure S.1).

**Figure S.1.**
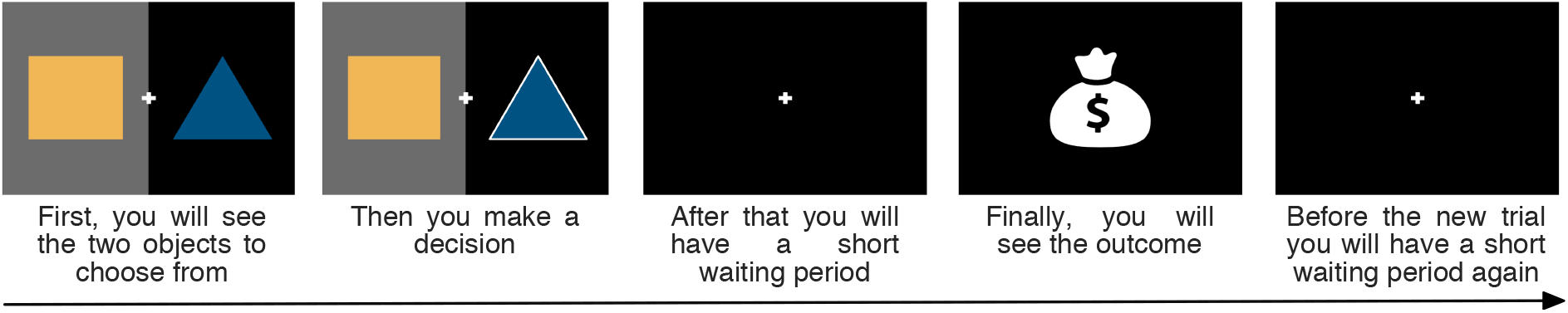
Participant instructions 1. The figure shows the sequence of events within a trial as presented to the participants in the instructions.

You will start the experiment with a balance of 0 points and any wins or losses will be registered to your account. After the experiment, in addition to your standard payment for participation, you will receive up to $30 depending on your final account. Note that your balance cannot get below 0 and if you do not earn additional money on the task, you will not be penalized and you will still receive the standard payment for your participation. We would however encourage you to try to earn as much as possible on the task. After each run we will show you your balance. A run consists of 80 trials, which takes about 20 minutes to complete. You will have two runs in the scanner.

As mentioned above, one of the objects is profitable and it will bring you a win most of the times and every now and then it will bring you loss. At the same time, the other object is not profitable and it will bring you a loss most of the time and a win every now and then. You won’t explicitly know which object is the profitable one and which is the non-profitable and you will need to conclude it from the outcomes. But be aware! These roles can switch, which means that the previously profitable object becomes non-profitable and the previously non-profitable object becomes profitable. Such a switch will happen only 1-4 times in the entire run and you will have enough trials without a switch to conclude which object is the profitable one.

Keep in mind that even the currently profitable object can from time to time deliver a loss and a couple of negative outcomes does not necessarily mean that a switch occurred. Similarly, even the non-profitable object can from time to time deliver a win and a couple of positive outcomes does not necessarily mean that a switch occurred. You can however assume that a switch has happened if you feel the previously rewarding object started to give you more losses than wins and the previously non-rewarding object started to bring you more wins than losses.

Before you do the test run, there is one more important aspect to the task: On each trial, one of the objects will be presented to you in front of a black background while the other object will be in front of a grey background. If you choose the object on the black-side, you will see the outcome of your choice. However, if you choose the grey-side object, the outcome will remain hidden from you but it will be registered to your account (Supplementary Figure S.2).

**Figure S.2.**
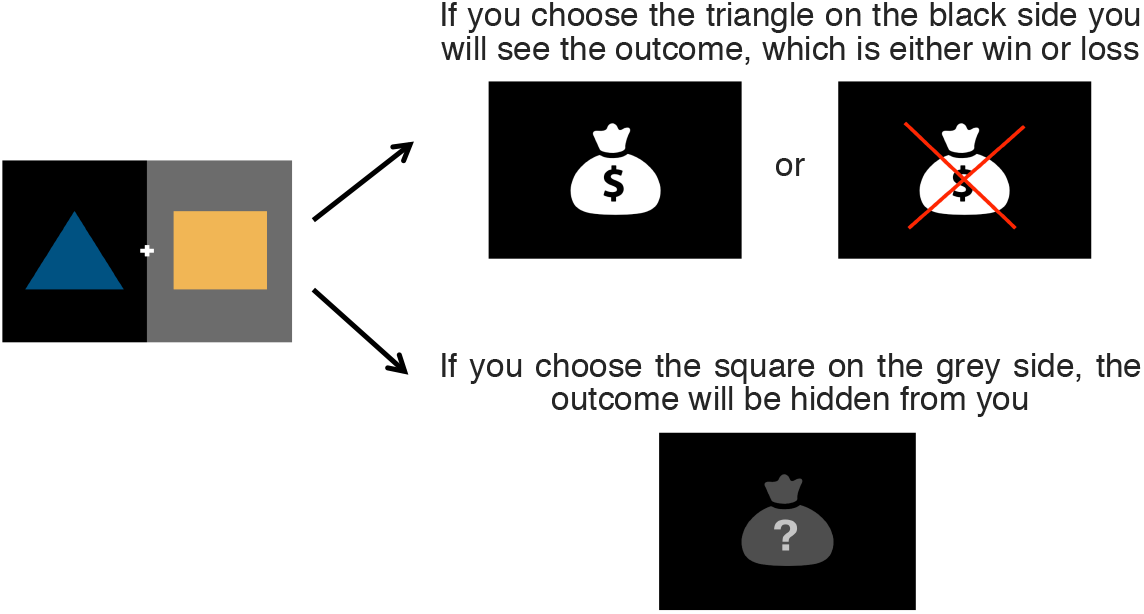
Participant instructions 2. The figure depicts the lucrativeness and informativeness associated with the actions as presented to the participants in the instructions.

You will now complete a test run, which will be just like the ones you will complete in the scanner. We will discuss all your questions to make sure you feel comfortable with the task before going in the scanner.

Note that from the perspective of the participants the specification of the available actions was doubly over-specified: on the one hand, the square and the triangle were also colored orange and blue, and on the other hand, one side of the screen was also indicated by a black background, while the other side was indicated by a gray background. For efficiency, in the main text, we only retain the the notions of squares and triangles for available rewarded actions, black and grey backgrounds for informative and non-informative actions, and we use colors to indicate the currently lucrative (yellow) and detrimental (blue) actions.

### S.3. Experimental state sequence

Table S.2 displays the definition of the state evolution function *f*. The upper table displays the state sequence of the first experimental run, the lower table displays the state sequence of the second run. *t* indexes the trial number and *s_t_* is the trial state.

**Table S.2.** Experimental state sequence.

|  |  |  |  |  |  |  |  |  |  |  |
| --- | --- | --- | --- | --- | --- | --- | --- | --- | --- | --- |
| $t$ | 1 | 2 | 3 | 4 | 5 | 6 | 7 | 8 | 9 | 10 |
| $s_t$ | (1, 1) | (1, 1) | (1, 2) | (1, 1) | (1, 2) | (1, 1) | (1, 1) | (1, 2) | (1, 1) | (1, 1) |
| $t$ | 11 | 12 | 13 | 14 | 15 | 16 | 17 | 18 | 19 | 20 |
| $s_t$ | (1, 1) | (1, 1) | (1, 2) | (1, 2) | (1, 2) | (1, 1) | (1, 2) | (1, 2) | (1, 2) | (2, 1) |
| $t$ | 21 | 22 | 23 | 24 | 25 | 26 | 27 | 28 | 29 | 30 |
| $s_t$ | (2, 1) | (2, 2) | (2, 1) | (2, 2) | (2, 1) | (2, 2) | (2, 2) | (2, 1) | (2, 2) | (2, 2) |
| $t$ | 31 | 32 | 33 | 34 | 35 | 36 | 37 | 38 | 39 | 40 |
| $s_t$ | (2, 1) | (2, 1) | (2, 2) | (2, 1) | (2, 2) | (2, 1) | (2, 2) | (2, 1) | (2, 2) | (2, 1) |
| $t$ | 41 | 42 | 43 | 44 | 45 | 46 | 47 | 48 | 49 | 50 |
| $s_t$ | (2, 2) | (2, 2) | (1, 2) | (1, 2) | (1, 1) | (1, 1) | (1, 2) | (1, 2) | (1, 2) | (1, 2) |
| $t$ | 51 | 52 | 53 | 54 | 55 | 56 | 57 | 58 | 59 | 60 |
| $s_t$ | (1, 1) | (1, 2) | (1, 1) | (1, 2) | (1, 2) | (1, 1) | (1, 1) | (1, 2) | (1, 1) | (1, 1) |
| $t$ | 61 | 62 | 63 | 64 | 65 | 66 | 67 | 68 | 69 | 70 |
| $s_t$ | (1, 1) | (1, 1) | (1, 2) | (2, 1) | (2, 1) | (2, 2) | (2, 1) | (2, 2) | (2, 1) | (2, 1) |
| $t$ | 71 | 72 | 73 | 74 | 75 | 76 | 77 | 78 | 79 | 80 |
| $s_t$ | (2, 2) | (2, 1) | (2, 1) | (2, 1) | (2, 2) | (2, 2) | (2, 2) | (2, 2) | (2, 2) | (2, 1) |
| $t$ | 1 | 2 | 3 | 4 | 5 | 6 | 7 | 8 | 9 | 10 |
| $s_t$ | (2, 1) | (2, 1) | (2, 1) | (2, 2) | (2, 1) | (2, 1) | (2, 1) | (2, 1) | (2, 2) | (2, 2) |
| $t$ | 11 | 12 | 13 | 14 | 15 | 16 | 17 | 18 | 19 | 20 |
| $s_t$ | (2, 1) | (2, 1) | (2, 2) | (2, 2) | (2, 1) | (2, 2) | (2, 2) | (2, 2) | (2, 2) | (2, 1) |
| $t$ | 21 | 22 | 23 | 24 | 25 | 26 | 27 | 28 | 29 | 30 |
| $s_t$ | (2, 1) | (2, 1) | (2, 2) | (1, 1) | (1, 2) | (1, 2) | (1, 2) | (1, 1) | (1, 2) | (1, 1) |
| $t$ | 31 | 32 | 33 | 34 | 35 | 36 | 37 | 38 | 39 | 40 |
| $s_t$ | (1, 2) | (1, 2) | (1, 2) | (1, 2) | (1, 2) | (1, 1) | (1, 1) | (1, 2) | (1, 2) | (1, 2) |
| $t$ | 41 | 42 | 43 | 44 | 45 | 46 | 47 | 48 | 49 | 50 |
| $s_t$ | (1, 2) | (1, 1) | (2, 2) | (2, 2) | (2, 2) | (2, 1) | (2, 2) | (2, 2) | (2, 2) | (2, 1) |
| $t$ | 51 | 52 | 53 | 54 | 55 | 56 | 57 | 58 | 59 | 60 |
| $s_t$ | (2, 2) | (2, 1) | (2, 2) | (2, 1) | (2, 1) | (2, 2) | (2, 1) | (2, 2) | (2, 1) | (1, 2) |
| $t$ | 61 | 62 | 63 | 64 | 65 | 66 | 67 | 68 | 69 | 70 |
| $s_t$ | (1, 1) | (1, 1) | (1, 2) | (1, 1) | (1, 1) | (1, 2) | (1, 1) | (1, 1) | (1, 1) | (1, 1) |
| $t$ | 71 | 72 | 73 | 74 | 75 | 76 | 77 | 78 | 79 | 80 |
| $s_t$ | (1, 2) | (1, 1) | (1, 1) | (1, 2) | (1, 1) | (1, 2) | (1, 1) | (1, 1) | (1, 1) | (1, 2) |

### S.4. Belief state, posterior predictive distribution, and KL-divergence

#### Belief state

Based on the probability distributions 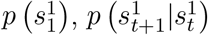 and 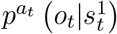, the agent’s belief state on trials *t* = 2, …, *T* can be recursively evaluated according to eq. (15). To show the validity of this equation, we first express the belief state as

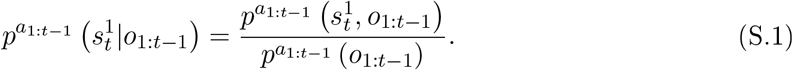

The numerator of eq. (S.1) can then be rewritten as

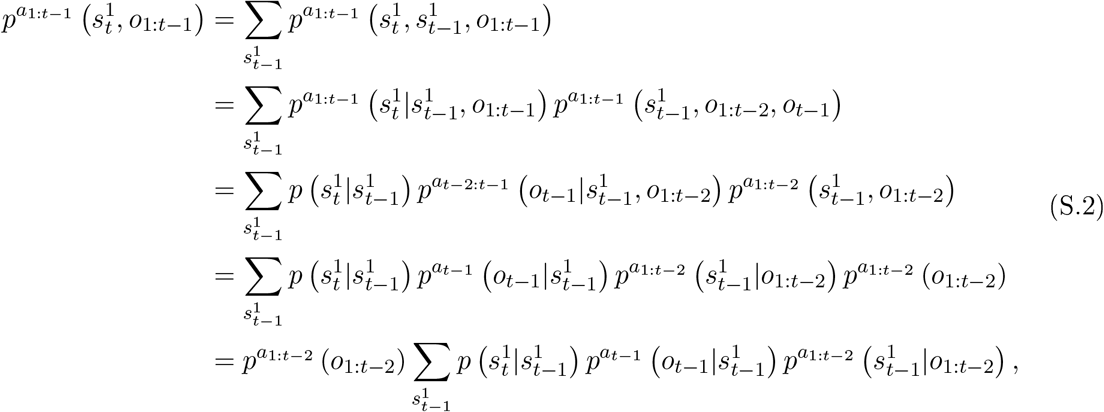

where we used the conditional independence of 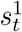 of *o*_1:*t*−1_ given 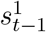 in the third equality and the conditional independence of *o_t−_*_1_ of all other random variables given 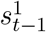 in the fourth equality.

Similarly, we can rewrite the denominator of eq. (S.1) as

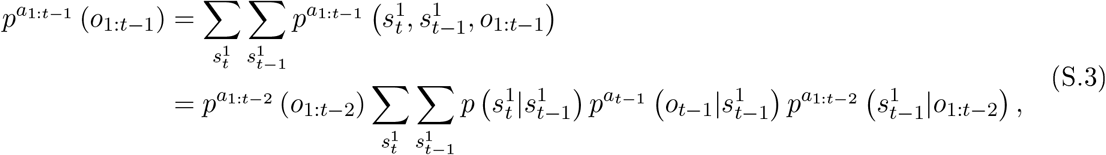

where in the last equality we used the numerator’s form derived in eq. (S.2). Finally, by substitution of (S.2) and (S.3) in (S.1), we obtain the belief state update formula of eq. (15) as follows:

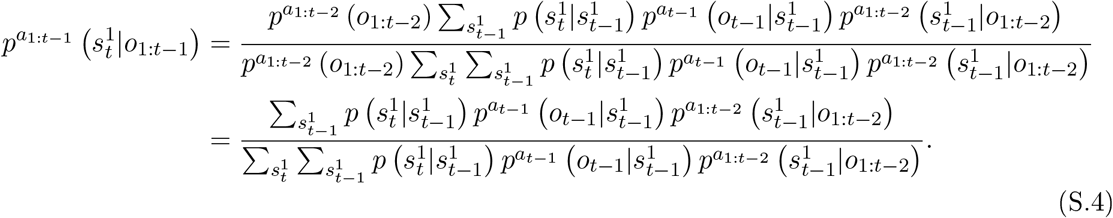

#### Posterior predictive distribution

Given the agent’s belief state 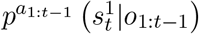 and the action-dependent state-conditional observation distribution 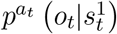, the posterior predictive distribution can be evaluated according to eq. (23). A proof of this equation is as follows:

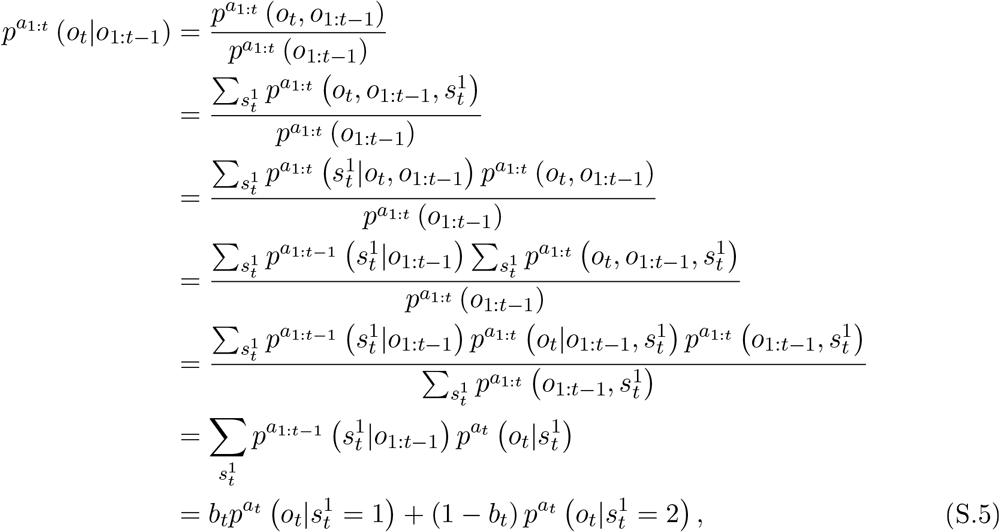

where in the last equality we substituted the belief state with its scalar representation.

#### KL-divergence

Recall that the KL-divergence for two distributions *p* and *q* of a discrete random variable *x* is defined as (Kullback & Leibler, 1951)

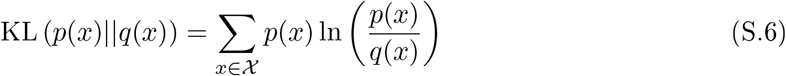

With

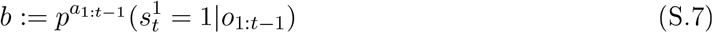

and thus

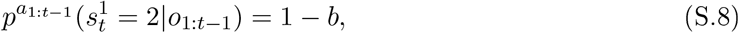

as well as

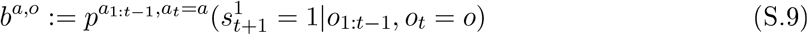

and thus

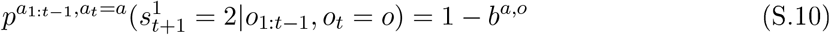

we have

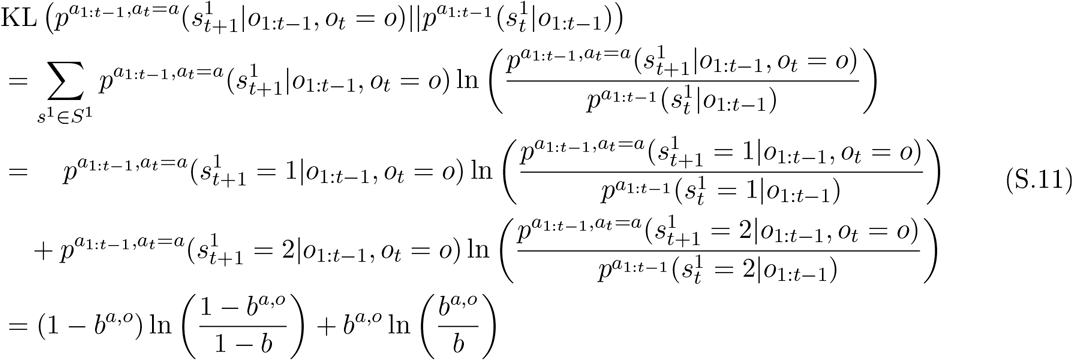

### S.5. Belief state and posterior predictive distribution implementation

For a concise implementation of the belief state update and the posterior predictive distribution formulas, we represent the agent’s probability distributions by stochastic vectors and stochastic matrices. Specifically, in the implementation of the agent model components as defined in *irb_modcomp.m*,

- 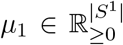 represents the initial belief state 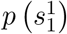. The *i*th entry of *μ*_1_ corresponds to the agent’s subjective uncertainty that the non-observable state component takes on value 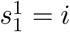 at trial *t* = 1. Formally,

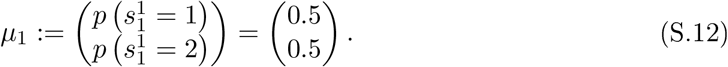
- 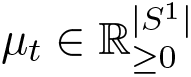 represents the belief state 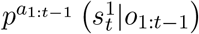 at trial *t*. The *i*th entry of *μ_t_* corresponds to the agent’s subjective uncertainty that the non-observable state component takes on value 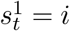 at trial *t* given the history of observations *o*_1:*t*−1_ and actions *a*_1:*t*−1_. Formally,

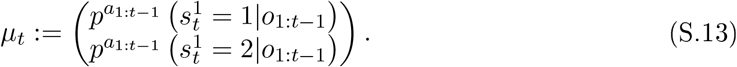
- 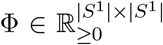 represents the state-state transition distribution 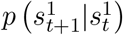. The *j*th entry of the *i*th row of Φ corresponds to the agent’s subjective uncertainty that the non-observable state component takes on the value 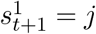 in trial *t* + 1 given that 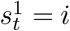 in trial *t*. Formally,

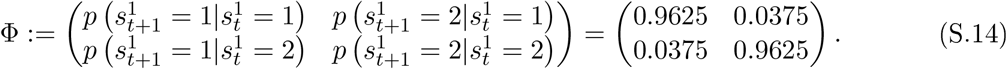
- 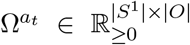 represents the action-dependent state-conditional observation distribution 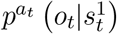 for action *a* ∈ *A*. The *k*th entry of the *i*th row of 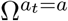 corresponds to the agent’s subjective uncertainty that the observation takes on the value *o_t_* = *k* given that the non-observable state component takes on the value 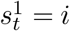 and the action value is *a_t_* = *a*. Formally, for the informative actions

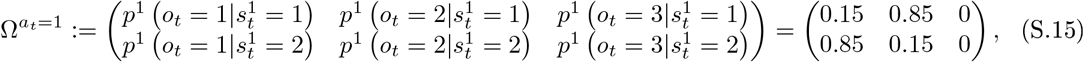

and

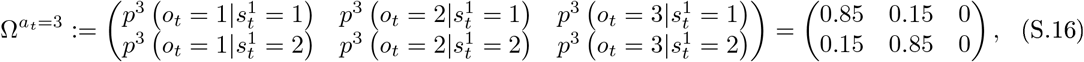

and for the non-informative actions

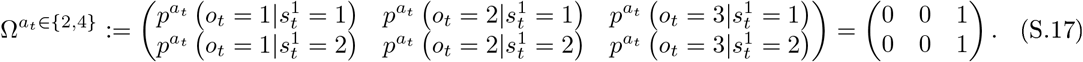
- 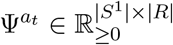 represents the action-dependent state-conditional reward distribution 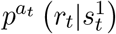 for action *a* ∈ *A*. The *l*th entry of the *i*th row of 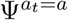 corresponds to the agent’s subjective uncertainty that the reward takes on the value *r_t_* = *l* − *m* given that the non-observable state component takes on the value 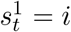 and action the value *a_t_* = *a*. Note that *m* is introduced to convert the linear indices to reward values and takes on the value 2 if *l* = 1 and the value 1 if *l* = 2. Formally,

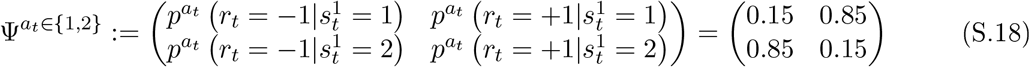

represents the action-dependent state-conditional reward distribution for the actions of choosing the square, and

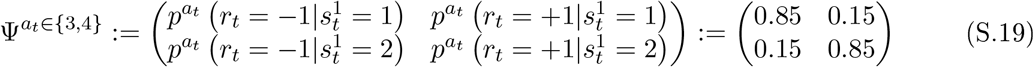

represents the action-dependent state-conditional reward distribution for the actions of choosing the triangle. Accordingly, 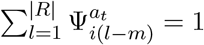.

Based on the definitions above and using the standard matrix product · as well as the element-wise (Hadamard) matrix product ◦, the agent’s belief state at trial *t* (eq. (15))can be written as

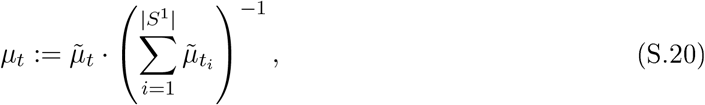

where

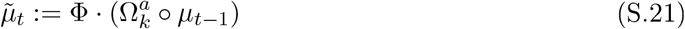

is the unnormalized belief state following action *a_t−_*_1_ = *a* and observation *o_t−_*_1_ = *k* and 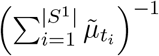 is the normalization constant. Here, 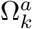 denotes the *k*th column of Ω^*a*^ and *μ*_*t*−1_ denotes the prior belief state on trial *t* − 1, which corresponds to eq. (S.13), if *t* − 1 > 1, and to eq. (S.12), if *t* − 1 = 1. Similarly, the posterior predictive distribution of agent model A2 (cf. eq. (23)) for action *a_t_* = *a* can be written as

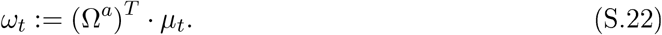

Prof. Dr. Dirk Ostwald

Abteilung Methodenlehre I

Institut für Psychologie

Otto-von-Guericke-Universitä t Magdeburg

Prof. Dr. Scott Brown

Editor in Chief

Computational Brain & Behavior

**Revised manuscript COBB-D-20-00031 “Human belief state-based exploration and exploitation in an information-selective symmetric reversal bandit task”**

Dear Prof. Dr. Scott Brown,

as encouraged by your letter of May 3rd 2021, please find attached our revised manuscript. We were delighted by your positive assessment of our initial revision and are thankful for your remaining suggestions to further enhance the clarity and potential impact of our work.

In line with your comments, we have (1) streamlined the manuscript’s outline to reflect the logical flow of the work, (2) added an explanatory overview of our agent-based behavioral modeling approach, and (3) documented the agent model codes in glossary form. Please find our detailed responses to your comments in our Response to Review letter below.

We believe that these changes have further improved the manuscript’s limpidity and we hope that you will find our revised manuscript to be suitable for publication in *Computational Brain & Behavior*.

Sincerely,

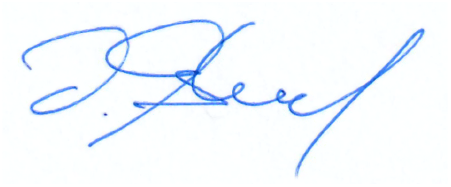

Prof. Dr. Dirk Ostwald

**Reviewer #1**

1. The manuscript is quite heavy on notation and formalism. This was something noted in the review reports on the earlier version. I appreciate the need for the notation, to provide clarity. However, it will also limit the impact that your manuscript can have, by confusing some sections of the audience. Your response letter makes good arguments for keeping the notation and layout largely the same, but I would like to suggest a relatively minor change. At the moment, after the empirical section ends the theory section begins very abruptly (line 227, section “Model Formulation”). For readers who are less familiar with formal math, the several pages of definition that follow will obscure the high-level (and interesting!) theoretical contributions. I suggest adding a small section under the “Model Formulation” heading, before the formalism, which provides a plain-English overview of the theoretical work to follow. This overview could explain the objectives of the modelling, and provide intuition about how the models explain the key aspects of the data. With an overview like this (or more, if you think so) readers will find the formalisms easier to understand and contextualise.

We thank you for this comment and have included the following paragraph at the beginning of the Agent-based behavioral modeling section:

2. The model names are not informative, and add cognitive burden for the reader. Instead of making the reader remember what C1, A3, etc. mean, is it possible to replace the codes with more psychologically meaningful names? Or, at least, have a table somewhere which reminds the reader of the key aspects of each model, like a glossary.

We thank you for this comment. We have added the table shown below to the revised Model Formulation section to document the key characteristics of each agent in glossary form.

**Table 1:**
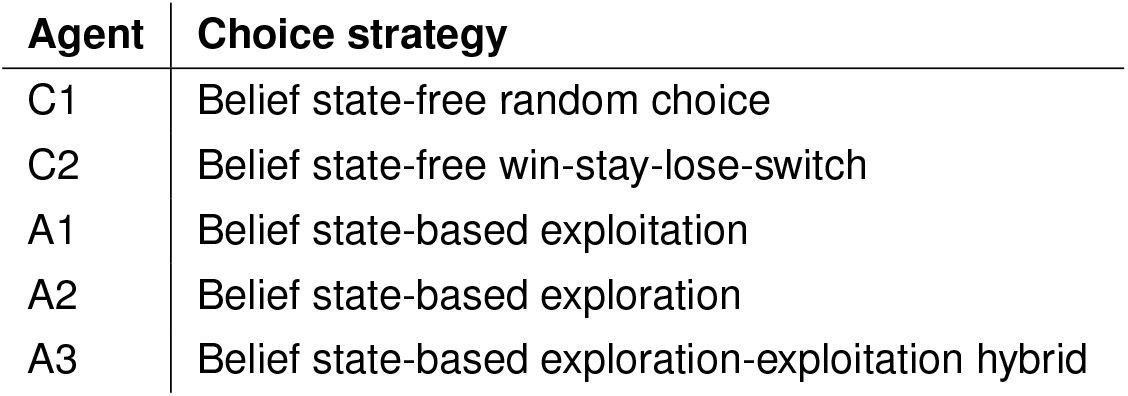
Agent model space. Agent model denominations (left column) and keywords highlighting central aspects of the respective agent’s choice strategy (right column).

We would prefer not to replace the agent denomination in general, because in our experience this comes at the risk of over-stating the generality of models developed. For example, if we were to rename agent model “A1” as “Belief state-based exploitative agent” this denomination would imply a scope that is not warranted by our formulation and implementation: there are many different probabilistic models that can implement belief states and there are many different ways in which belief states can be used for exploitative choices. The generic denomination as “A1” thus emphasizes the (necessary) idiosyncrasy of the current model space.

3. I realise you have already tried several re-orderings of the sections. However, I agree with some earlier sentiments that the behavioural results would be more sensibly set out earlier. This journal does not require APA format, and does not mandate any particular names or order of sections. If you think a different order would be better, please use that. One that comes to mind for me is: Introduction; Methods; Behavioural Results; Model Formulation; Model Results; Discussion.

We thank you for this comment and are happy to deviate from APA format in favor of an outline that better suits the content of our work. In line with your suggestion, the principle outline of the manuscript is now as follows: *Introduction; Experimental methods; Descriptive analyses; Agent-based behavioral modeling; Discussion*. Notably, the section *Descriptive analyses* now includes both the descriptive data-analytical methods and the results of these analyses, strongly enhancing the readability of this section. Similarly, the section *Agent-based behavioral modeling* includes the formal model development, the methods used for assessing the models in light of the experimental data, the results of our model and parameter recovery analyses, as well as the final model comparison and post-hoc model validation results. As for *Descriptive analyses*, the logical flow of this section is now much enhanced.

